# Integrin alpha1 beta1 promotes interstitial fibrosis in a mouse model of polycystic kidney disease

**DOI:** 10.1101/2024.10.18.619080

**Authors:** C. Grenier, I-H. Lin, DJM. Peters, A. Pozzi, R. Lennon, RW. Naylor

**Affiliations:** Manchester Cell-Matrix Centre, Division of Cell-Matrix Biology and Regenerative Medicine, School of Biological Sciences, Faculty of Biology Medicine and Health, The University of Manchester, Manchester, United Kingdom; Bioinformatics Core Facility, Faculty of Biology Medicine and Health, University of Manchester, Manchester, United Kingdom; Department of Human Genetics, Leiden University Medical Center, Leiden, Netherlands; Department of Medicine, Division of Nephrology and Hypertension; Department of Veterans Affairs, Nashville, Tennessee, USA

## Abstract

Fibrosis is the cause of end-stage kidney failure in patients with Autosomal Dominant Polycystic Kidney Disease (ADPKD). The molecular and cellular mechanisms involved in fibrosis are complex and anti-fibrotic therapies have so far failed to make an impact on patient welfare. Using unbiased proteomics analysis on the *Pkd1*^nl/nl^ mouse, we found that expression of the integrin α1 subunit is increased in this model of ADPKD. In human ADPKD tissue and two single cell RNA kidney disease datasets, *ITGA1* was also upregulated. To investigate the functional role of this integrin subunit in ADPKD, we generated a *Pkd1*^nl/nl^*Itga1*^−/−^ mouse. We observed a significant reduction in kidney volume and kidney dysfunction in mice lacking the integrin α1 subunit. Kidneys from *Pkd1*^nl/nl^*Itga1*^−/−^ mice had smaller cysts and reduced interstitial expansion and tubular atrophy. Picrosirius red staining identified a restriction in collagen staining in the interstitium and the myofibroblast marker α smooth muscle actin was also downregulated. Myofibroblast cell proliferation was reduced in *Pkd1*^nl/nl^*Itga1*^−/−^ mice and primary fibroblast cultures demonstrated an abrogated fibrogenic phenotype in integrin α1-depleted fibroblasts. These results highlight a previously unrecognised role for the integrin α1 subunit in kidney fibrosis.

## Introduction

Polycystic kidney disease (PKD) affects ∼1 in 1000 individuals[1] and accounts for 15% of chronic kidney disease (CKD) cases. Autosomal dominant polycystic kidney disease (ADPKD) is the most common subtype and is mostly caused by variants in the *PKD1* and *PKD2* genes. *PKD1* and *PKD2* encode for polycystin-1 (PC1) and PC2, respectively[2–4] and the PC1-PC2 complex forms a transient receptor potential (TRP) ion channel that has many roles in cell function[5], most of which are poorly understood. The major feature of PKD is bilateral formation of kidney cysts, which induce chronic inflammation and fibrotic scarring. Combined, these pathologies lead to kidney failure in ∼50% of individuals with ADPKD by age 60 years.

The common pathological feature and final manifestation of CKD is interstitial fibrosis. Fibrosis is characterised by the replacement of functional tissue with a collagen-rich extracellular matrix (ECM)[6], and there are currently no effective anti-fibrosis treatments. In CKD, the pathological indication of fibrosis involves interstitial expansion, leukocyte invasion, basement membrane thickening, tubular atrophy, vascular rarefaction, and fibrogenesis (accumulation of fibrillar ECM). In ADPKD, fibrotic scarring and cyst formation are coupled; preventing fibrosis reduces cyst formation[7], and precluding cyst formation reduces inflammatory signals that initiate fibrosis[8].

In fibrosis, stromal cells in the interstitium are activated by inflammatory signals, such as TGFβ, and transdifferentiate to proto-myofibroblasts, and then senescent myofibroblasts[9]. Myofibroblasts are the main source of new ECM in fibrotic kidneys and are predominantly derived from resident fibroblasts and pericytes[10]. Inhibition of mesenchymal cell proliferation and myofibroblast differentiation, ECM deposition and contractility is likely to restrict fibrosis in CKD. Cell-ECM signalling is integral to these processes as the interaction of cells with their surrounding microenvironment determines their fibrogenic potential[11, 12]. Integrins are the principal cellular transmembrane receptors that bind ECM, and they exist as heterodimers of one alpha and one beta subunit. In mammals there are 18 alpha subunits and 8 beta subunits which form 24 distinct receptors with specific ECM ligand binding activity[13]. Integrins connect extracellular ligands with the cytoskeleton and form a signalling nexus called a focal adhesion[14]. The collagen-binding integrin alpha subunits (*ITGA1*, *ITGA2*, *ITGA10*, *ITGA11*) all dimerise with integrin beta 1. Genetic knockouts of these integrin alpha subunits in mice are generally well tolerated[15–17]. Genetic deletion of the integrin α1 subunit increases glomerulosclerosis and glomerular deposition of collagen IV in response to adriamycin-induced glomerular injury[18] or a streptozotocin model of diabetic nephropathy[19]. In the unilateral ureter obstruction (UUO) model of kidney fibrosis, loss of integrin α1 caused an increase in tubulointerstitial fibrosis[20]. In the skin, *Itga1*^−/−^ mice also display increased collagen production and an irregular wound response[21]. In dermal fibroblasts, integrin α1 positively regulates cell proliferation through activation of the adaptor protein Shc[22]. A pro-proliferative function for integrin α1 is supported in cancer studies, where its inhibition attenuates cell proliferation and metastasis[23, 24]. These results suggest that integrin α1 is an anti-fibrotic and pro-proliferative receptor. However, in other contexts depletion of integrin α1 may reduce fibrosis. A triple knockout of the three major collagen binding receptors *Itga1*, *Itga2* and *Itga11* is non-lethal, and fibrosis is attenuated in these mice[25], suggesting that collagen-binding integrins positively control the development of fibrosis. In Alport syndrome, loss of integrin α1 is protective[26], and this may be in part due to reduced interstitial fibrosis[27]. A recent study showed that Alport mice crossed onto the *Itga1*^−/−^ mice treated with the lowering blood pressure drug Ramipril had increased lifespan and delayed fibrosis compared to Alport mice[28]. These results suggest the role of integrin α1 in regulating fibrotic responses may be disease-context and cell-context specific.

Using mass spectrometry-based proteomics, we found upregulation of integrin α1 in a hypomorphic mouse model of ADPKD (*Pkd1*^nl/nl^). In parallel, we found integrin α1 localisation in fibroblasts and myofibroblasts in human healthy and ADPKD kidney tissue. To test the biological relevance of increased integrin α1 in ADPKD, we performed global genetic depletion of *Itga1* in *Pkd1*^nl/nl^ mice and observed a striking reduction in cyst size and interstitial fibrosis. These findings were associated with reduced interstitial fibrosis, fewer interstitial macrophages and a reduction in fibrogenic markers in primary fibroblast cultures taken from *Pkd1*^nl/nl^*Itga1*^−/−^ kidneys. Overall, our findings indicate that integrin α1 positively regulates the development of cysts and fibrogenesis in ADPKD and therefore represents a new therapeutic target.

## Methods and Materials

### Animal husbandry and maintenance

All mouse handling, breeding and disposal was performed in adherence to the culture of care policies of the University of Manchester. The ARRIVE guidelines 2.0 were adhered to when designing, implementing and recording experimental outcomes to ensure best practice and reliable results. Urine samples were collected for urinary Albumin Creatinine Ratio (uACR) measurements by transferring individual mice to a separate sterile cage. Mice were left for a maximum of 15 minutes and spots of urine were collected in this timeframe (at least 50 µl was collected). Urine samples were stored at −20°C and when ready were processed using a Creatinine Parameter Assay kit (R&D systems, cat# KGE005) and a mouse Albumin ELISA kit (Bethyl Laboratories, cat# E99-134). Kidneys for proteomics analysis were collected at P1 and P28, when body weight and two kidney weight measurements were made immediately after schedule 1 killing. For P28 mice, blood samples were collected by cardiac puncture and left overnight at 4°C to coagulate. Serum was collected after spinning samples down at 1200 g for 5 minutes at 4°C to pellet the blood cells. Serum was stored at - 80°C. Urea Nitrogen measurements were made on serum samples using the Blood Urea Nitrogen (BUN) Colorimetric Detection kit (Arbor Assays, cat# K024-H1). Kidneys to be used in proteomics and Western blot analyses were flash frozen in liquid nitrogen and stored at −80°C. Kidney tissue to be used in qPCR analyses was placed in TRIzol reagent and stored at −80°C.

Sex as a biological variable was not relevant to this study given no sex-specific differences have been observed in the *Pkd1*^nl/nl^ mouse at the stages analysed. However, we did analyse equal numbers of male and female mice to ensure no variance was observed between sexes. No differences were detected and so the analyses shown are of both sexes combined.

### Histology and Immunofluorescence

For light microscopy, kidneys were fixed in 4% PFA for 3 days at 4°C, paraffin embedded in a tissue processor and then mounted. 5 µm sections were cut on a Leica RM2255 microtome and slides were left to dry overnight at 37°C. Haemotoxylin and eosin (H&E) and picrosirius red (PSR) staining was performed on a Leica Autostainer XL CV5030. Images of whole kidneys were collected as tiled images on an Olympus BX63 upright microscope using a 20x / 0.75 UApo/340 objective and captured and white-balanced using a DP80 camera (Olympus) in colour mode through CellSens Dimension v1.16 (Olympus). Images were then processed and analysed using Fiji ImageJ (http://imagej.net/Fiji/Downloads) (see statistical analyses section below for quantification processing of collected images). For immunofluorescence analyses, dried 5 µm FFPE sections were also used. Slides were dewaxed as follows; 5 mins Xylene, 5 mins Xylene, 2 mins 100% Ethanol, 2 mins 100% Ethanol, 2 mins 90% Ethanol, 2 mins 75% Ethanol, then placed in distilled water. Primary antibodies used were integrin α1 (Cell Signalling Technologies 15574T, 1:200, Citrate antigen retrieval), plectin (Boster Bio PB9430, 1:250, Citrate antigen retrieval), αSMA (ThermoFisher 14-9760-82, 1:250, citrate antigen retrieval, used for images in Figure S8), αSMA-Cy3 (Merck C6198 1:500 (added with secondaries) no antigen retrieval needed but can be used on citrate or TrisHCl treated tissue), PCNA (Merck P8825, 1:200, TrisHCl antigen retrieval), F4/80 (Proteintech 81668-1-RR, 1:250, TrisHCl antigen retrieval). Species-specific secondary antibodies used were AlexaFluor488, AlexaFluor594, AlexaFluor647 (ThermoFisher, all at 1:500 dilution). Following dewaxing, antigen retrieval was performed. For the citrate pH 6.0 and TrisHCl pH 9.0 antigen retrieval methods, slides were placed in pre-warmed antigen retrieval solution and incubated in a 70°C water-bath for 30 mins, then removed and allowed to cool to room temperature for an additional 30 mins. After antigen retrieval slides were washed 3x 5 mins in PBS and 0.05% Tween20 (PBSTw), then blocked for 2 hours in PBS containing 2% bovine serum albumin, 1% donkey serum, 0.5% Triton X-100. Primary antibodies were added, and slides were left overnight at 4°C. Slides were washed 3x 15 mins with PBSTw and then secondary antibodies and Hoechst (5 µg/ml) nuclear stain were added for 1 hour at room temperature in the dark. Slides were then washed 3x 5 mins in PBSTw and left to dry in the dark for 30 mins before being mounted in ProLong Gold Antifade Mountant and cover slipped. Images were collected on an Olympus BX63 upright microscope using a 20x / 0.75 UApo/340 objective and captured and white-balanced using an DP80 camera (Olympus) in monochrome mode through CellSens Dimension v1.16 (Olympus). Specific band pass filter sets for DAPI, FITC, Cy3, Texas red and Cy5 were used to prevent bleed through from one channel to the next. Images were then processed and analysed using Fiji ImageJ (http://imagej.net/Fiji/Downloads). For single plane confocal images shown in Figure S2B, images were collected on a Leica TCS SP8 AOBS inverted confocal using a 63*x* / 1.4 NA objective. The confocal settings were as follows, pinhole 1 airy unit, scan speed 1000Hz unidirectional, format 1024 x 1024. Images were collected using hybrid detectors with the following detection mirror settings; FITC 494-530nm using the white light laser with 488nm (20%) and DAPI 405 (10%) laser lines respectively. When it was not possible to eliminate crosstalk between channels, the images were collected sequentially. The super-resolution images in Figure 3C were collected on a Zeiss LSM 880 Airyscan confocal microscope using a 63x / 1.4 Plan-Apochromat objective. The confocal settings were as follows, pinhole [1 airy unit], scan speed 400Hz unidirectional, format 1024 x 1024. Images were collected using an Airyscan detector with the following spectral channel settings; FITC 494-530nm; Texas red 602-665nm, using the 488nm (20%) and 561nm (100%) laser lines respectively. When acquiring 3D optical stacks, the confocal software was used to determine the optimal number of Z sections. Only the maximum intensity projections of these 3D stacks are shown in the results.

### Proteomics analysis

Kidneys were taken from P1 or P28 mice, flash frozen in liquid nitrogen and stored at −80°C. For label-free quantitative proteomics, samples were then prepared using the S-trap protocol as described in [29]. Samples were first chemically fractionated to enrich kidneys for detection of extracellular matrix (ECM) proteins[30]. After capsule removal, samples were cut into half and one half was minced into 1 mm^3^ pieces in 500 µl of sterile PBS. Half kidney fragments were then passed through a 100 µm sieve and washed through with 500 µl PBS. The final volume of 1 ml of suspended kidney tissue was centrifuged at 14,000g for 5 mins at 4°C. After supernatant removal, kidney weights were obtained. Kidney fragments from each replicate was then re-suspended in a normalised volume of extraction buffer (10 mM Tris, 150 mM NaCl, 1% Triton X-100, 25 mM EDTA, Roche protease inhibitor) such that 1 mg kidney/µl solution. 50 µl of each normalised kidney mass was then passed to fresh tubes for fractionation. Samples were left for 30 mins at 4°C with mild vortex every 10 minutes. They were then centrifuged at 14,000g for 5 mins at 4°C. The supernatant was collected (fraction 1) and the pellet was re-suspended in 50 µl alkaline detergent buffer (20 mM NH_4_OH and 0.5% Triton X-100 in PBS). The remaining sample was incubated for 30 mins at 4°C with mild vortex every 10 minutes and then centrifuged again at 14,000g for 5 mins at 4°C. The supernatant was removed (fraction 2), then samples were re-suspended in 25 µl of 1X S-trap buffer (5% SDS, 50 mM TEAB pH 7.5) and heated at 70°C for 10 mins. Samples were then transferred to Covaris glass tubes for sonication on Covaris LE220+ (Parameters: duration 500s, peak power 500, duty factor 40%, cycles/burst 50, average power 200). Sonicated samples (now in solution) were transferred to fresh tubes (fraction 3). For the soluble fraction, 8.3 µl of fraction 1 and 16.7 µl fraction 2 were combined with 25 µl of 2X S-trap lysis buffer. For the less soluble fraction, all 25 µl of fraction 3 was used for preparation for the mass spectrometer. Following these steps, we performed reduction (DTT) and alkylation (IAM), then in-column trypsin digestion, peptide elution and R3 desalt as described in [29].

Sample analysis was performed by liquid chromatography-tandem mass spectrometry (LC-MS/MS). For this study, we used an Orbitrap Fusion Lumos Tribid Mass Spectrometer, sample runtime for each replicate was 60 mins. Raw spectra data were acquired and analysed with MaxQuant[31]. Statistical analysis was performed in the Perseus platform[32]. Results were filtered for significant FDR master proteins identified with ≥1 unique peptide detected in ¾ biological replicates.

### Single-cell RNA-seq analysis

The human FACS sorted CD10^-^ single-cell RNA-seq (scRNA-seq) data by[10] was obtained from the Zenodo data archive (https://zenodo.org/records/4059315). Downstream analysis and visualisation of the data was performed in R environment (v4.3) following the workflow documented in Orchestrating Single-Cell Analysis with Bioconductor[33].

A SingleCellExperiment class object was created using the provided UMI count data, gene-level and cell-level annotations. The log-normalized expression values were computed using the “multiBatchNorm” function from the batchelor R package (v1.18.1). The per-gene variance of the log-expression profile was modelled using the “modelGeneVarByPoisson” function from the scran R package (v1.30.2) and top 2500 highly variable genes were selected. The mutual nearest neighbours (MNN) approach implemented by the “fastMNN” function from the batchelor R package was used to perform batch correction.

The first 50 dimensions of the MNN low-dimensional corrected coordinates for all cells were used as input to produce the t-stochastic neighbour embedding (t-SNE) projection and uniform manifold approximation and projection (UMAP) using the “runTSNE” and “runUMAP” functions from the scater R package (v1.30.1) respectively.

### Kidney Tissue Atlas scRNA-seq Data

The scRNA-seq data in the h5Seurat format was retrieved from the Kidney Tissue Atlas of the Kidney Precision Medicine Project (https://atlas.kpmp.org/).

The dataset was loaded and analysed in R. It has log-normalised expression data from a total of 77 individuals, including 26 healthy participants (43,380 cells), 14 acute kidney injury (AKI) patients (51,167 cells) and 37 chronic kidney disease (CKD) patients (130,630 cells). A dotplot was created to show the RNA expression of *ITGA1* across 54 cell types in the three enrolment categories. Cell type abbreviations: aFIB, Adaptive / Maladaptive / Repairing Fibroblast; aPT, Adaptive / Maladaptive / Repairing Proximal Tubule Epithelial Cell; aTAL1, Adaptive / Maladaptive / Repairing Thick Ascending Limb Cell; aTAL2, Adaptive / Maladaptive / Repairing Thick Ascending Limb Cell; ATL, Ascending Thin Limb Cell; B, B Cell; CCD-PC, Cortical Collecting Duct Principal Cell; cDC, Classical Dendritic Cell; CNT, Connecting Tubule Cell; CNT-IC-A, Connecting Tubule Intercalated Cell Type A; CNT-PC, Connecting Tubule Principal Cell; C-TAL, Cortical Thick Ascending Limb Cell; cycEPI, Cycling Epithelial Cell; cycMNP, Cycling Mononuclear Phagocyte; cycT, Cycling T Cell; DCT1, Distal Convoluted Tubule Cell Type 1; DCT2, Distal Convoluted Tubule Cell Type 2; dFIB, Degenerative Fibroblast; dPC, Degenerative Principal cell; dPT, Degenerative Proximal Tubule Epithelial Cell; DTL1, Descending Thin Limb Cell Type 1; dVSMC, Degenerative Vascular Smooth Muscle Cell; EC-AEA, Afferent / Efferent Arteriole Endothelial Cell; EC-AVR, Ascending Vasa Recta Endothelial Cell; EC-GC, Glomerular Capillary Endothelial Cell; EC-LYM, Lymphatic Endothelial Cell; EC-PTC, Peritubular Capillary Endothelial Cell; FIB, Fibroblast; IC-A, Intercalated Cell Type A; IC-B, Intercalated Cell Type B; IMCD, Inner Medullary Collecting Duct Cell; MAC-M2, M2 Macrophage; MAST, Mast Cell; MC, Mesangial Cell; MDC, Monocyte-derived Cell; MON, Monocyte; M-TAL, Medullary Thick Ascending Limb Cell; MYOF, Myofibroblast; ncMON, Non-Classical Monocyte; NK1, Natural Killer 1; NKT, Natural Killer T Cell; PC, Principal Cell; pDC, Plasmacytoid Dendritic Cell; PEC, Parietal Epithelial Cell; PL, Plasma Cell; POD, Podocyte; PT-S1/S2, Proximal Tubule Epithelial Cell Segment 1&2; PT-S3, Proximal Tubule Epithelial Cell Segment 3; REN, Renin-positive Juxtaglomerular Granular Cell; T, T Cell; T-CYT, Cytotoxic T Cell; tPC-IC, Transitional Principal-Intercalated Cell; VSMC, Vascular Smooth Muscle Cell; VSMC/P, Vascular Smooth Muscle Cell / Pericyte.

### qRT-PCR

TRIzol suspended tissue was sonicated to lyse cells and reveal nucleic acids. RNA extraction was performed using the RNAeasy mini kit (Qiagen, cat# 74104). cDNA was diluted to 5 ng/µl and mixed with 7.8 µM of primers and added in equal volume to 2X Power SYBRTM Green PCR Master Mix (Thermofisher, cat# 4367659). Primers used are shown in the table below. All primers were validated by standard curves prior to use, with only primers having an R^2^>90 and PCR efficiency of 90-110% used for analysis.

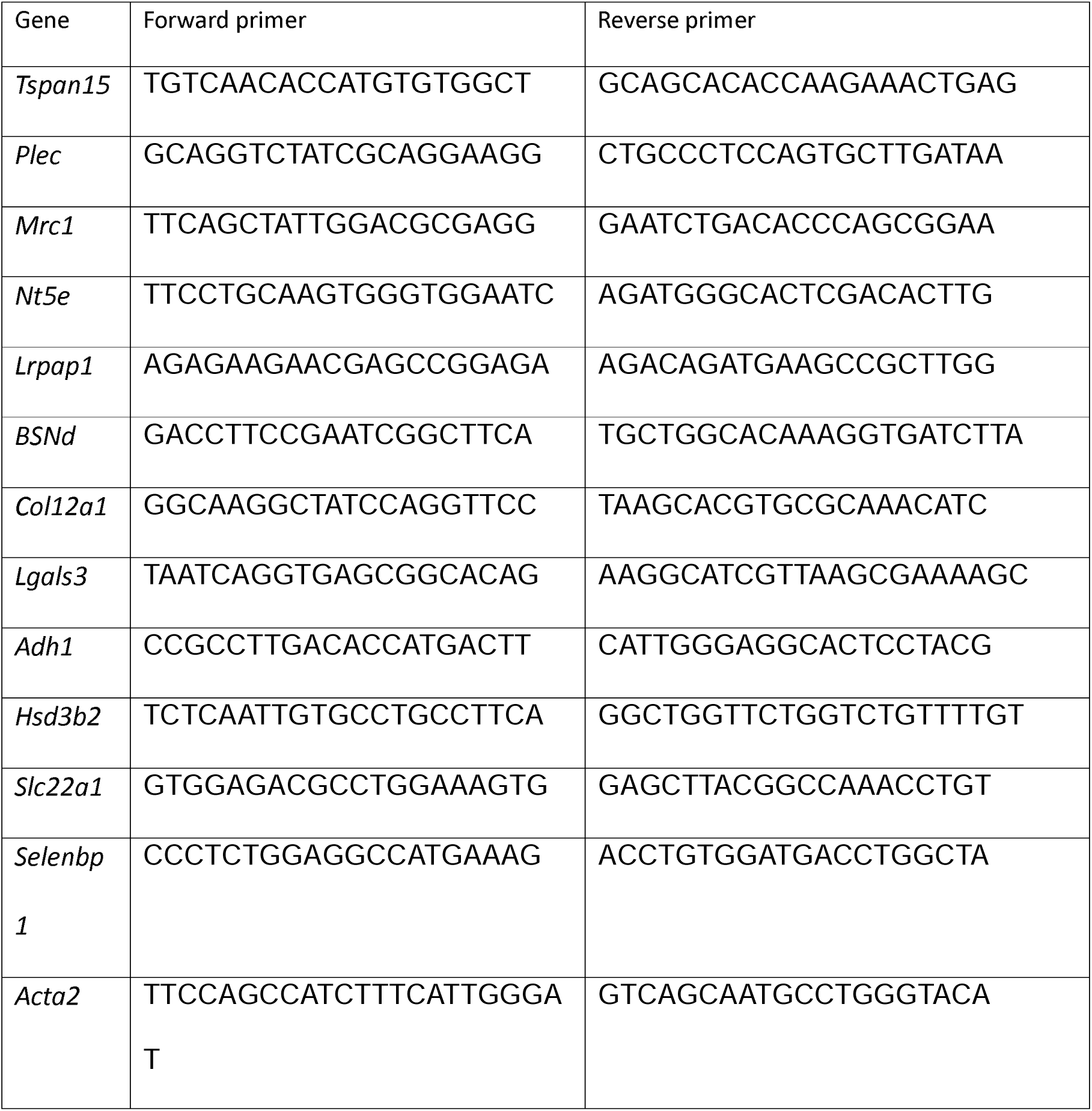
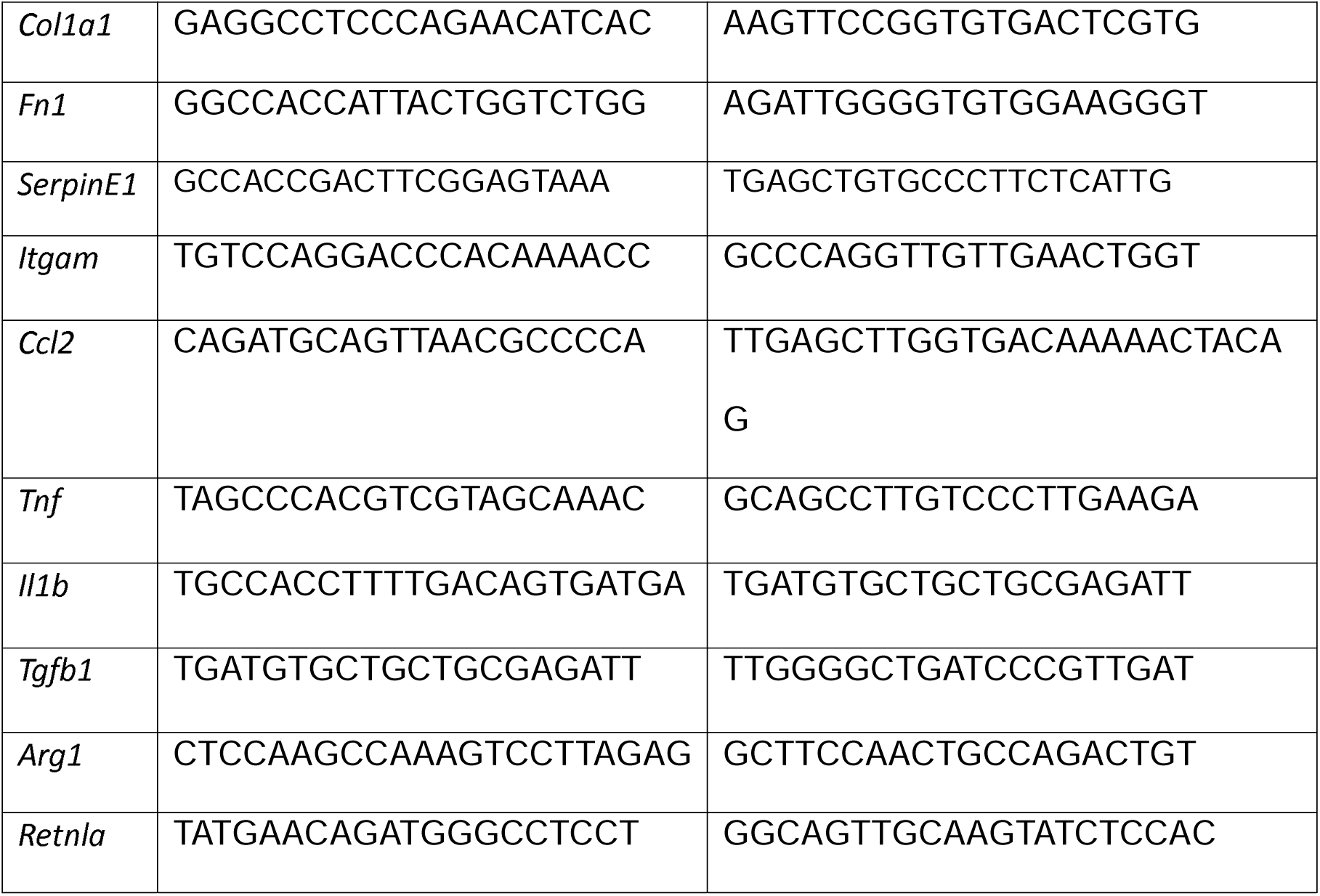

The PCR reaction was run on a Bio-Rad CFX96 Touch Real-Time PCR machine. All analyses of the data used the ΔΔCt method of quantification. RT-PCRs were run in triplicate with -RT and -cDNA controls. *Ppia* was used as the housekeeping gene for normalising Ct values based on the findings of [34].

### Western blot

Quarter kidneys stored at −80°C were placed in 200 µl RIPA buffer (150 mM NaCl, 10 mM Tris HCl, 1% Na deoxycholate, 0.1% SDS) with protease inhibitor (Merck, cat# 05892791001). To this, four 1.6 mm diameter stainless steel beads (Next Advance, cat# SSB16) were added and the tissue was beaten in a Bullet Blender Storm 24 2x 1 min. 50 µl of lysate was removed and placed in a fresh tube, with the remaining lysate stored at −20°C. 150 µl of RIPA buffer, 70 µl NuPAGE LDS Sample Buffer (ThermoFisher, cat# NP0007), and 15 µl 1M DTT were added to the 50 µl of isolated lysate, which was then heated at 95°C for 5 mins and stored at −20°C. For protein gels, 10 µl of reduced lysate was added to each well of a pre-cast gel with 4 µl of Precision Plus Protein standards loaded to determine molecular size. Protein gels were run at 200 Volts for 45 mins in 1X MES Running buffer, and then transferred to nitrocellulose membrane at 30 volts for 70 mins. Membranes were then blocked in Intercept (PBS) Blocking buffer (Li-Cor, cat# 927-70001) for 1 hour and incubated with primary antibodies overnight at 4°C in PBSTw. After 3x 5mins washes in PBSTw, secondary antibodies were added for one hour in MilliQ water. Membranes were imaged on a Li-Cor Odyssey CLx. Quantification of Western blots was performed in FIJI ImageJ (http://imagej.net/Fiji/Downloads).

### Primary fibroblast culture

Primary fibroblasts were cultured based on a modified method taken from [35]. Kidneys were isolated from mice and placed in chilled sterile PBS immediately. Kidneys were minced into 1 mm^3^ pieces in 1 ml digestion buffer (0.1% collagenase, 2.5 U/ml dispase II, 2 mM CaCl_2_, 10 mM HEPES, and 150 mM NaCl in water, filter sterilized). Minced kidney tissue from a whole kidney was transferred to 4 ml of digestion buffer in a sterile 50 ml tube and incubated at 37°C for 90 mins at 100 rpm. Digested tissue was then pushed through a 70 µm sieve into a fresh 50 ml tube. 20 ml of washing buffer (0.05 M EDTA in PBS, filter sterilised) was added and the cells were centrifuged at 260g for 5 mins to pellet. Supernatant was removed and 10 ml of DMEM/ 10% FBS/ 1% PenStrep fibroblast complete growth media was added, and the cells were centrifuged again at 260g for 5 mins to pellet. The cells were re-suspended in 10 ml fibroblast complete growth media and split to two 100 mm cell culture dishes. Cells were left for 3 days and then split (1:3) and left a further 3 days then split (1:3) to a 6-well plate and 8-well chamber slides (for EdU and cell spreading assays, 3×10^5^ cells per chamber). For EdU proliferation assay, cells were left to adhere for 24 hours then 1X EdU was added, and the cells were left for another 24 hours before fixation in 4% PFA overnight at 4°C. Cells were washed with PBS, then stained for EdU following the Click-iT EdU Cell Proliferation kit protocol manual (ThermoFisher, cat# C10340). Cells were then further stained for the nuclear marker Hoechst and αSMA. Proliferation rate was quantified by counting at least 300 nuclei and measuring the % of these that were EdU^+^. Fixed cells were also stained for phalloidin/αSMA to measure the number of primary fibroblasts that retained a myofibroblast state. For cell spreading assays, chambers of an 8-chamber slide were coated with either vitronectin, collagen I or collagen IV. 3×10^5^ cells were seeded onto coated surfaces (and uncoated controls) and left to adhere for four hours before being fixed in 4% PFA. Cells were visualised by staining for actin filaments; cells were washed 2x 5 mins in PBS containing 3% BSA, left in PBS containing 0.5% Triton X-100 for 20 mins, then incubated 1X Phalloidin-Alexa fluor 647 in PBS with 0.5% Triton X-100. Staining was left for one hour and then cells were washed with PBSTw 3x 5mins, their chamber box removed, left to dry for 30 mins, then mounted in Prolong Gold antifade mountant and cover slipped. Fluorescent imaging of stained primary cells was performed on an Olympus BX63 upright microscope. For contractility assays, we followed the protocol described by [36]. We performed NaOH titration for optimised collagen solidification for this protocol prior to cell treatments. Adherent fibroblasts were trypsinized and 300,000 cells were isolated to a 15 ml falcon tube in 1.2 ml of fibroblast complete growth media (DMEM/ 10% FBS/ 1% PenStrep). To this, we added 0.6 ml of 3 mg/ml stock collagen solution treated with pre-determined volume of 1 M NaOH and immediately mixed the cells and then transferred 0.5 ml to three separate wells of a 24-well plate. The cells were left for 30 mins at room temperature until the hydrogels had solidified and then the gels were dissociated from the wall of the wells using the tip of a 2 µl pipette tip. The cells were left for 48 hours, and the final diameter of the gel was measured on a Leica M205 FA upright Stereomicroscope using a 5x / 0.50 PlanAPO LWD objective and captured using a DFC 365FX (Leica) camera through LAS AF v3.1.0.8587software (Leica). Brightfield images were then processed and analysed using Fiji ImageJ (http://imagej.net/Fiji/Downloads).

### Statistical analyses

For PSR analysis on whole kidneys, the Olympus viewer plugin for Fiji was used to open .vsi files. The highest resolution was chosen in the Group Level window. When the image opened, the freehand selection tool was used to mark the area of the kidney, and the area was measured in µm^2^. The image was converted to 16-bit grayscale and threshold analysis was performed to mark the fibrotic area (measured in µm^2^). Given cystic kidneys had large areas of non-parenchymal space, we subtracted the cyst area from the total kidney area to get % fibrotic area (PSR^+^ collagen fibres area (µm^2^)/(total kidney area (µm^2^) - cystic area (µm^2^))). The cystic area was measured using particle analysis after thresholding, with only particles >200 µm^2^ measured as this was the size threshold by which we reliably detected the particle as a cyst rather than a tubule/duct lumen. This same process of measuring cystic index (% kidney area), average cyst size and cyst number was determined using H&E stained slides for the analysis given in Figure 5. For measurement of fibrotic index in fluorescent images, such as red fluorescent PSR (Figure 6B) and αSMA immunostained whole kidneys (Figure 1F and 1G, and Figure 6C), similar analysis was performed as described above, but a binary mask was generated in thresholding using the fluorescent signal rather than a colorimetric signal. For the analysis of human kidney samples, three randomly selected areas were imaged by confocal microscopy in each of the patient biopsies. These were then analysed using thresholding in FIJI to obtain % area coverage for integrin α1 and αSMA. For the PCNA/αSMA image analysis shown in Figure 7, whole kidney section views were analysed. Firstly, a binary mask was generated in the αSMA Cy3 channel (Process>Binary>Make Binary), which was then selected (Edit>Selection>Create Selection) and added to ROI list (Ctrl T). This ROI was then placed over the PCNA channel image and non-overlapping staining was deleted using the Edit>Clear Outside function. This image was then made into a binary mask using thresholding and PCNA^+^ nuclei in cells co-labelled with αSMA were measured in terms of number and area. Blind analysis was performed where possible, however we note that the phenotypes were easily discernible without quantification when comparing either wild-type and PKD kidneys, wild-type and PKD/*Itga1*^−/−^ kidneys and PKD and PKD/*Itga1*^−/−^ kidneys. All statistical tests were performed in GraphPad Prism 9.1.2; Grouped analyses – Two-way ANOVA (or mixed model). N numbers for each experiment are given in the text. Voronoi plots were generated in RStudio using logFC and Gene name to determine cell size. The PantherDB version 18.0 was used for pathway groupings. Only the top 100 upregulated proteins are shown, with the seed function set to 10.

**Figure 1:**
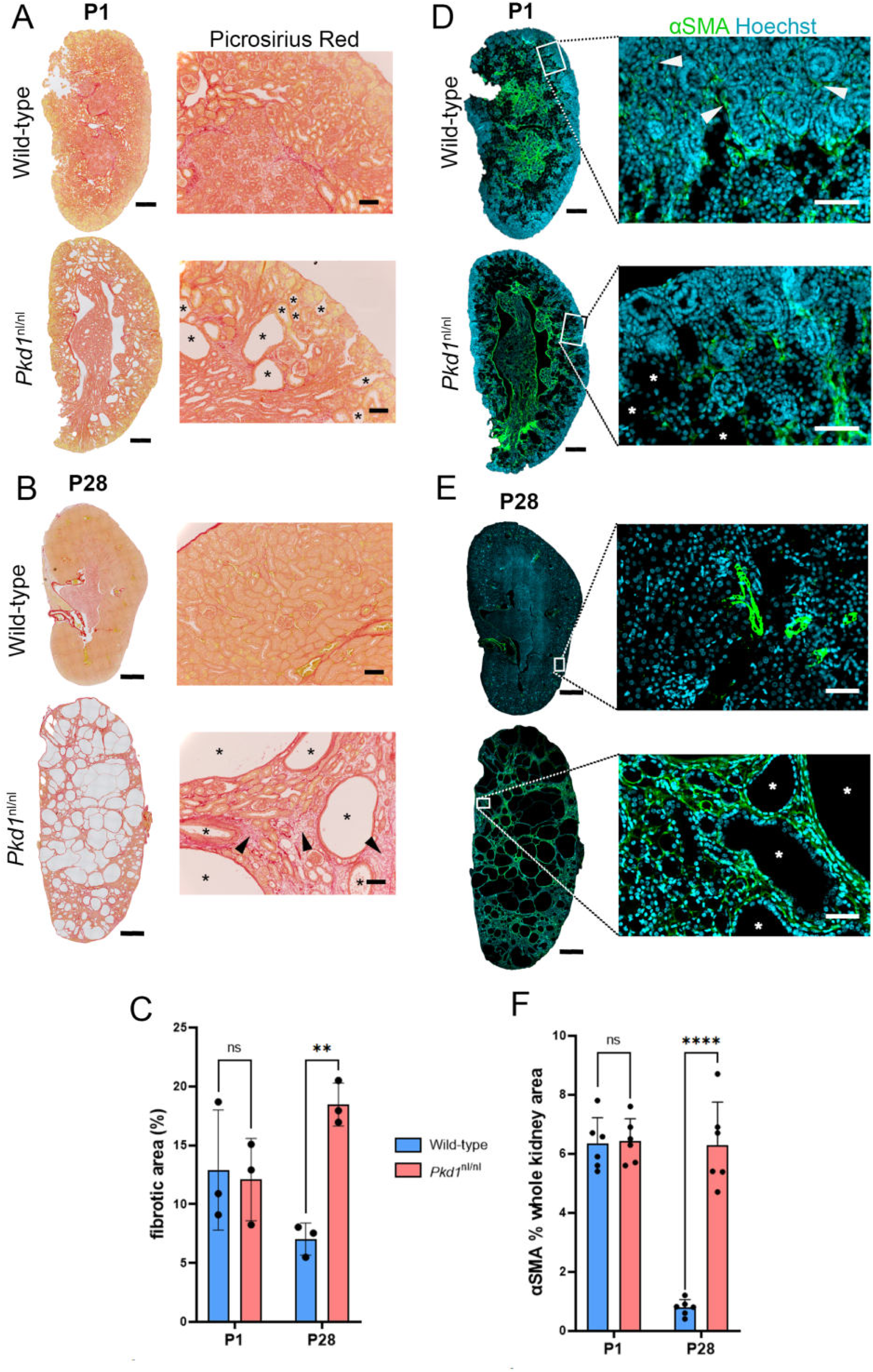
Interstitial fibrosis is established by P28 in Pkd1^nl/nl^ kidneys. A and B) Picrosirius Red (PSR) staining of P1 (A) and P28 (B) kidneys, whole kidney views shown on the left and higher magnification views in right panels. Arrowheads in B highlighting areas of collagen-rich fibrosis, asterisks indicate individual cysts. C) Histogram showing the % fibrotic area (measured from PSR staining) in wild-type and *Pkd1*^nl/nl^ kidneys at P1 and P28. D and E) Alpha smooth muscle actin (αSMA) staining of P1 (D) and P28 (E) kidneys, whole kidney views shown on the left and higher magnification views in right panels. Arrowheads in D indicate αSMA^+^ myofibroblasts in the nephrogenic zone of the P1 kidney cortex, asterisks indicate individual cysts. F) Histogram showing the % area of kidney that is αSMA in wild-type and *Pkd1*^nl/nl^ kidneys at P1 and P28. Scale bars: Whole kidney P1 views in A) and D) 200 μm, whole kidney P28 views in B) and E) 1000 μm, higher magnification panel views in A) and B) 100μm, higher magnification views in D) and E) 50 μm.

**Figure 2:**
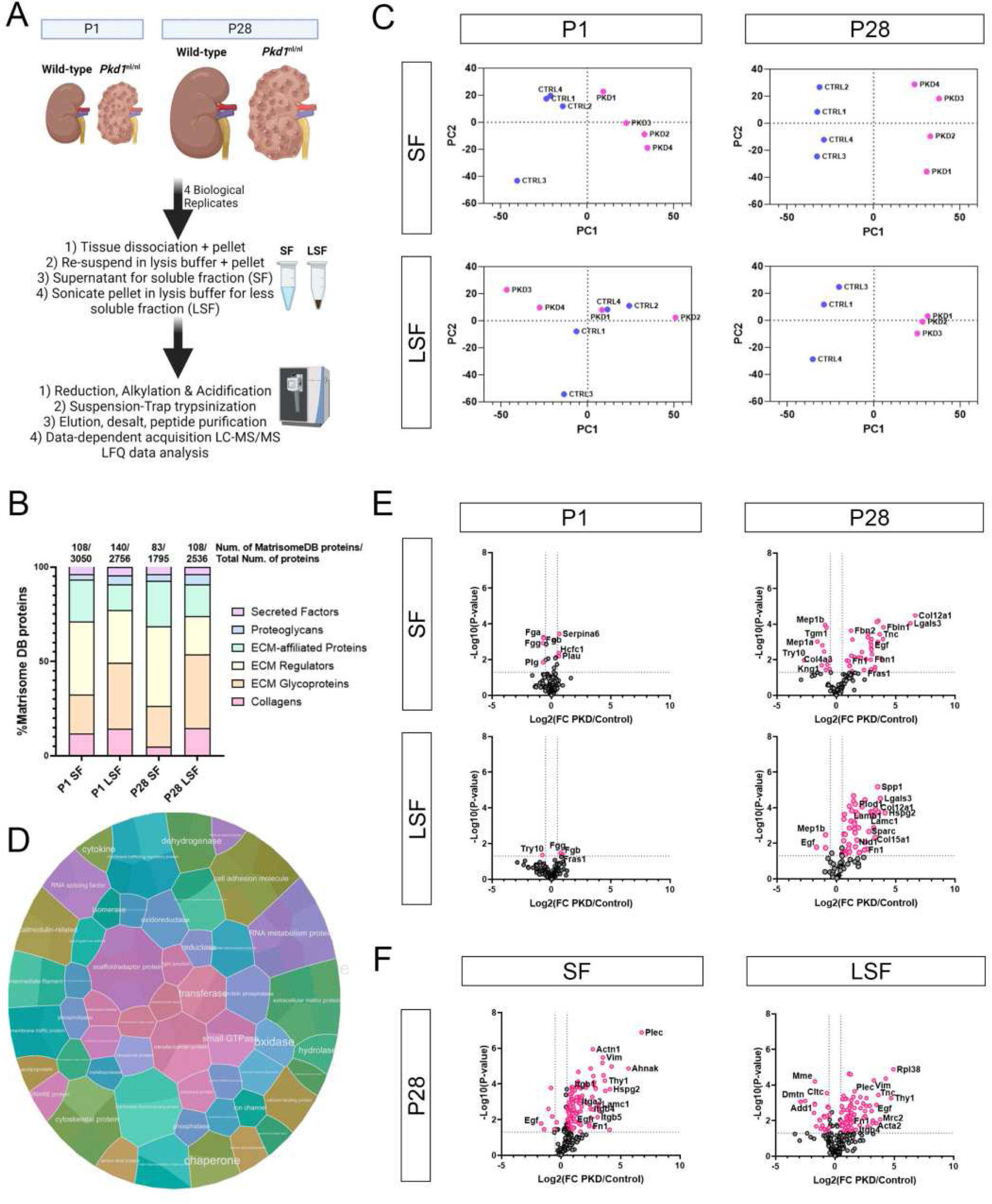
MS-based proteomics analysis identified significant changes in Pkd1^nl/nl^ adhesome at P28. A) Schematic of experimental plan and sample preparation for proteomics analysis. B) Histogram showing proportion and identification of ECM proteins in the datasets for each sample group. C) Principal component analysis for SF and LSF at P1 and P28. D) Voronoi plot showing Panther analyses for top 100 upregulated proteins in the P28 SF. E) Volcano plots showing the Log2 fold change in protein abundance of MatrisomeDB proteins comparing *Pkd1*^nl/nl^ (PKD) kidneys to *Pkd1*^+/+^ (Control) kidneys for the SF and LSF at P1 and P28. F) Volcano plots showing the Log2 fold change in protein abundance of focal adhesion proteins comparing *Pkd1*^nl/nl^ (PKD) kidneys to *Pkd1*^+/+^ (Control) kidneys for the SF and LSF at P28.

**Figure 3:**
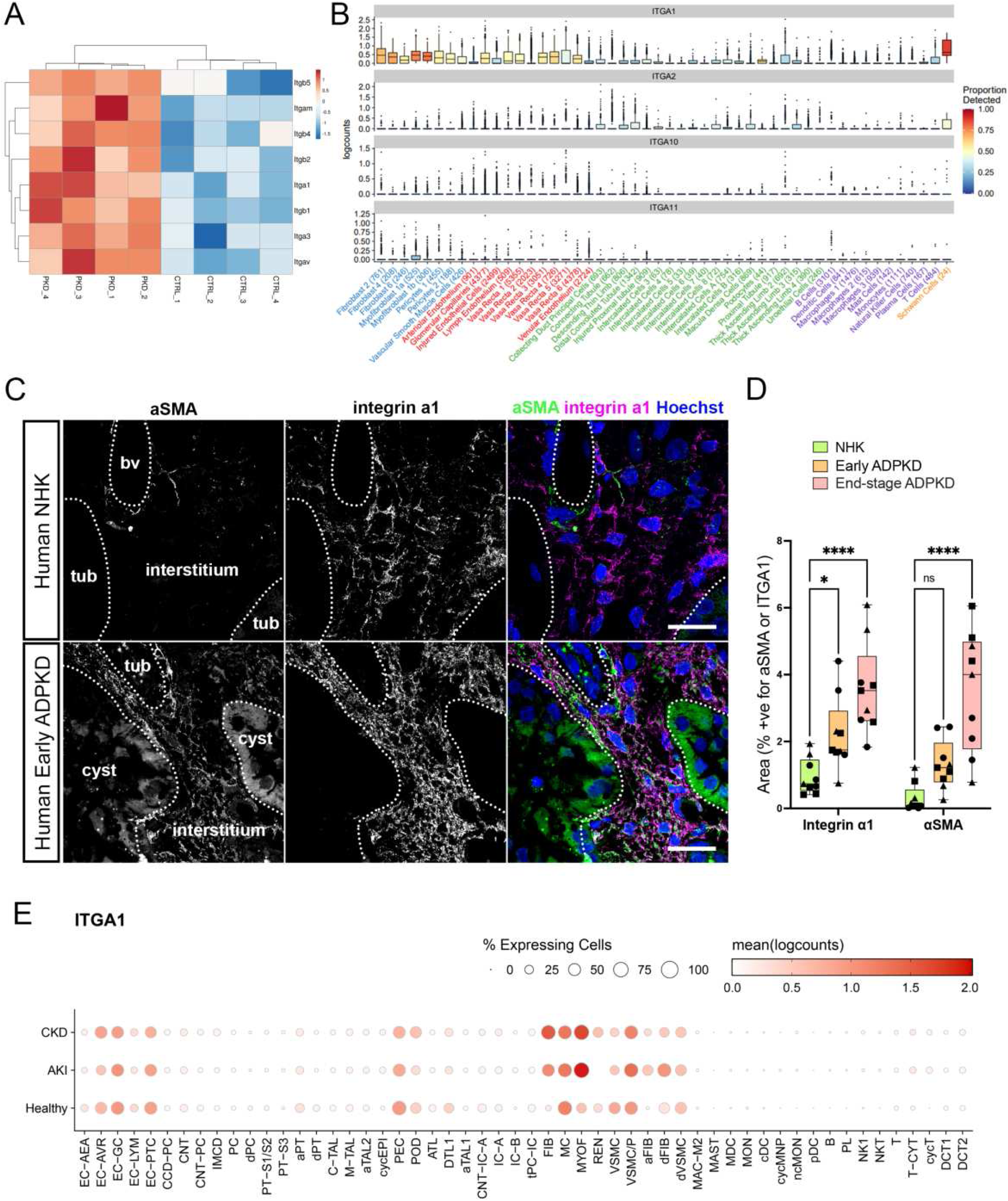
Integrin α1 is upregulated in Pkd1^nl/nl^ and human kidneys and is expressed in the interstitial mesenchyme. A) Heat map showing expression of all the integrin subunits detected in the *Pkd1*^nl/nl^ kidney proteomics dataset. B) Box and violin plots showing human kidney scRNA-seq data for *ITGA1*, *ITGA2*, *ITGA10* and *ITGA11*. C) Panels showing super-resolution microscopy images of human normal healthy kidney (top panels) and early stage ADPKD human kidney (bottom panels). Left panels are labelled for αSMA, middle panels are labelled for integrin α1 and right panels show the merge. Scale bar indicates 20 μm. D) Box and violin plots showing the extent of αSMA^+^ immunostaining (by area) in the indicated groupings. Each symbol represents a biological replicate, for each replicate three random snapshots were taken. E) Dot plot of *ITGA1* expression in a scRNA-seq dataset published by the kidney precision medicine project. For cell cluster abbreviation definitions, see Methods, MyoF, Myofibroblast; Fb, Fibroblast.

**Figure 4:**
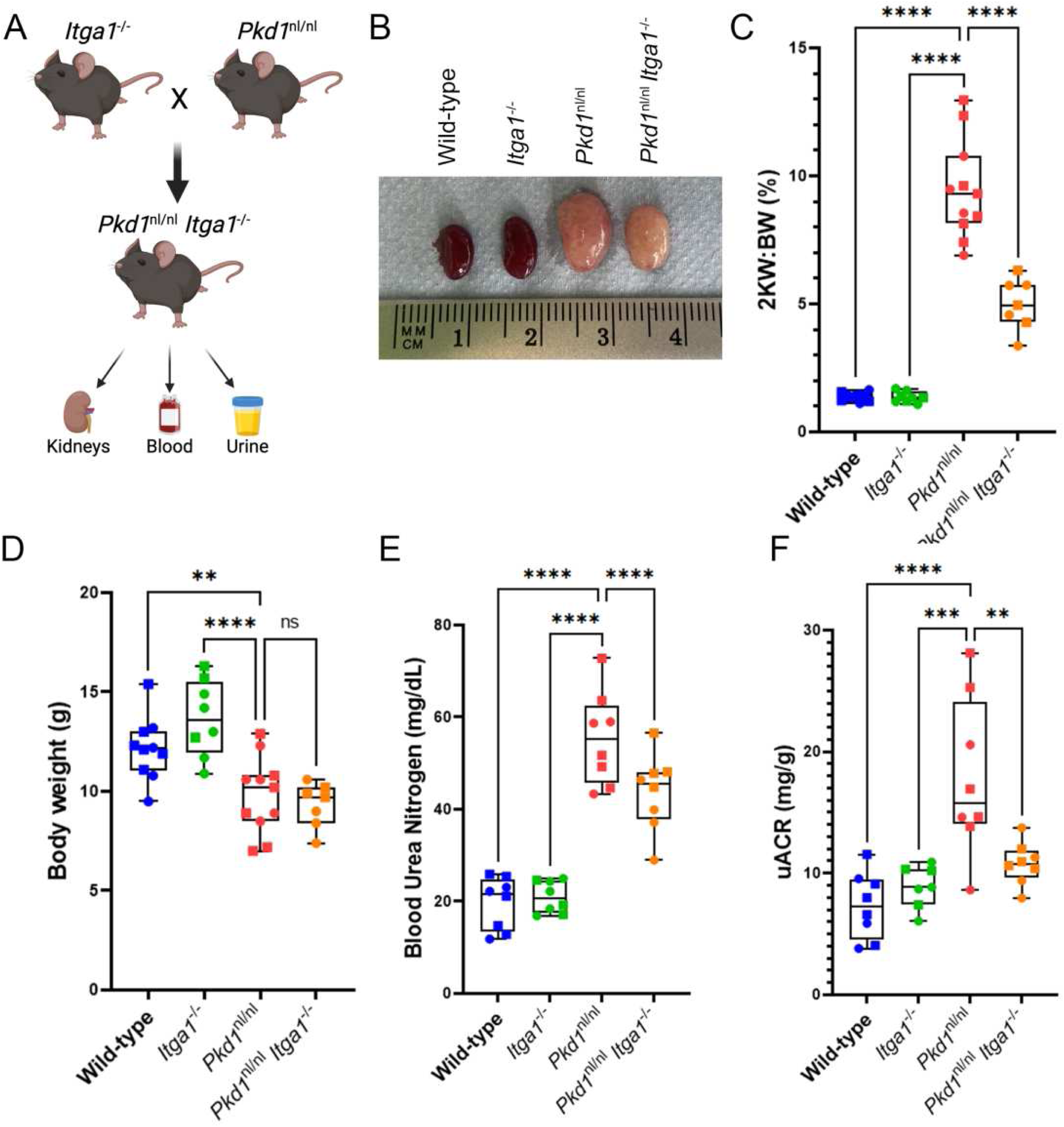
Integrin α1 depletion attenuated kidney pathology associated with Pkd1^nl/nl^ mice. A) Schematic showing experimental setup for mouse crossings and tissue collection. B) Photograph of fixed kidneys isolated from mice of indicated genotypes at P28. C)-F) Box and violin plots of % 2 kidney weight to body weight ratio (2KW:BW, C), body weight (grams, D), blood urea nitrogen (mg/dL, E) and urinary albumin to creatinine ratio (uACR, mg/g, F) of indicated genotypes. In dataplots in C)-F), square symbols identify males, circle symbols identify females.

**Figure 5:**
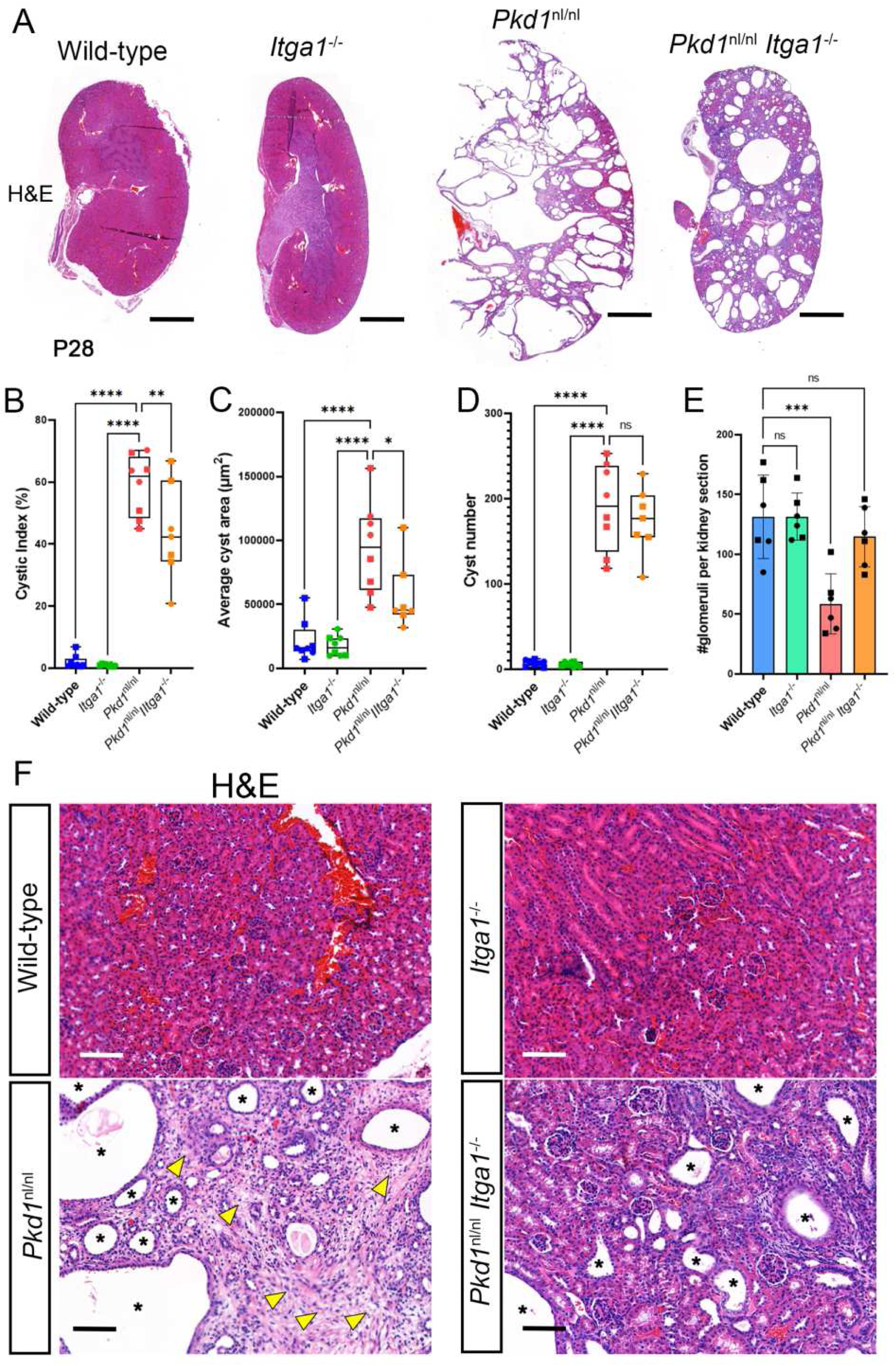
Cyst growth is reduced in Pkd1^nl/nl^ mice depleted of integrin α1. A) Haematoxylin and Eosin (H&E) stained whole P28 kidney views of indicated genotypes. Scale bars indicate 1000 μm. B)-E) Box and violin plots of cystic index (B), average cyst area (μm^2^, C), cyst number (D) and glomerular counts (E) of indicated genotypes. F) Panels showing higher magnification views of H&E-stained kidneys of indicated genotypes. Asterisks indicate cysts, arrowheads highlight areas of interstitial expansion and nephron atrophy. Scale bars indicate 100 μm. In dataplots shown in B)-E), square symbols identify males, circle symbols identify females.

**Figure 6:**
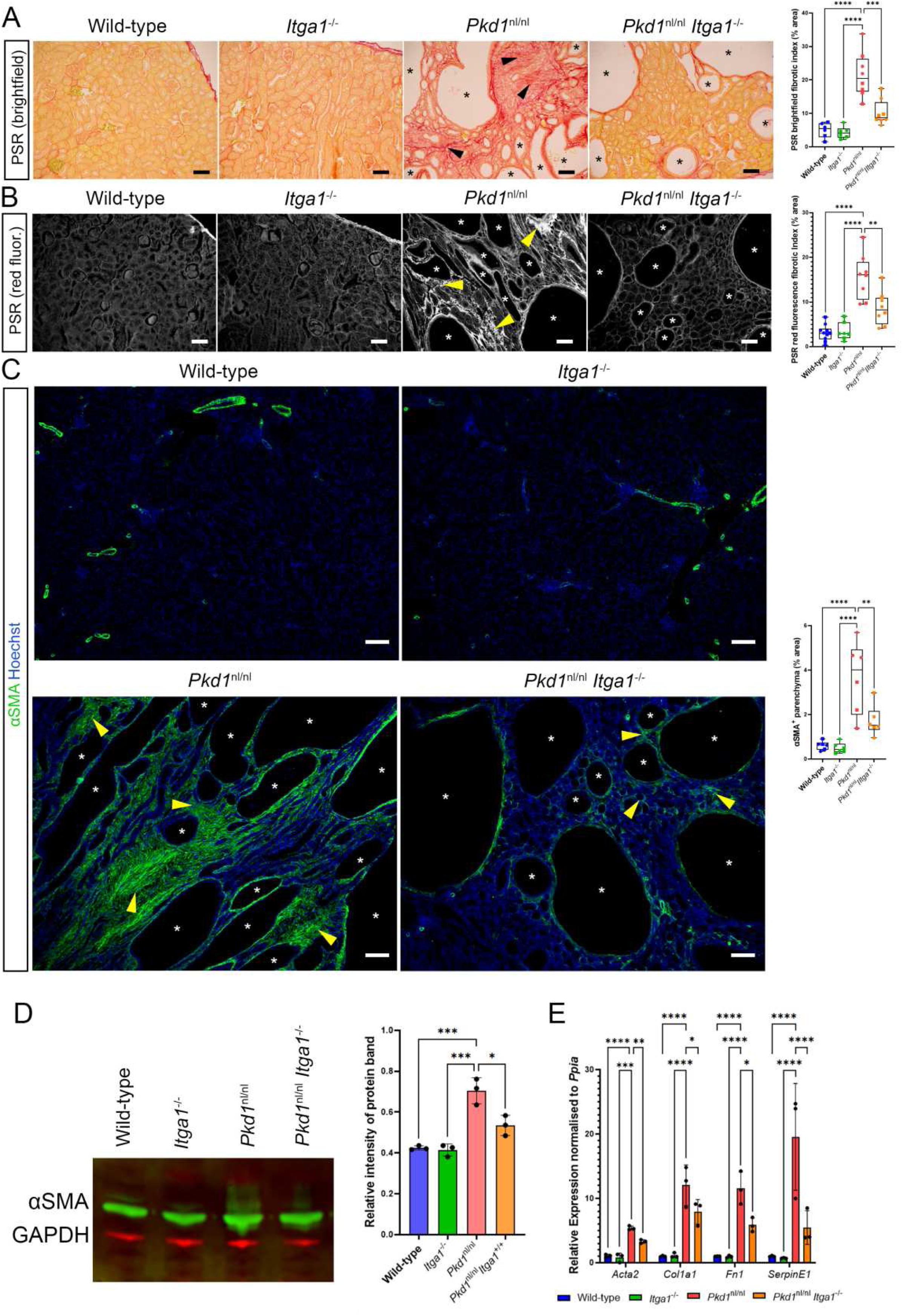
Integrin α1 depletion reduced interstitial fibrosis in Pkd1^nl/nl^ kidneys. A) Panels show brightfield views of Picrosirius red (PSR) staining on P28 kidneys of indicated genotypes. Black arrowheads show areas of extensive interstitial fibrosis, asterisks indicate individual cysts. Box and violin plot on the right shows’ quantification of the fibrotic area index for the indicated genotypes. Scale bar indicates 50 μm. B) Panels show red channel fluorescence views of Picrosirius red (PSR) staining on P28 kidneys of indicated genotypes. Yellow arrowheads show areas of extensive interstitial fibrosis, asterisks indicate individual cysts. Box and violin plot on the right shows’ quantification of the fibrotic area index for the indicated genotypes. Scale bar indicates 50 μm. C) Panels show αSMA immunostaining (green) and Hoechst staining (blue) on P28 kidneys of indicated genotypes. Arrowheads show areas of extensive αSMA^+^ labelling (areas of extensive fibrosis), asterisks indicate individual cysts. Box and violin plot on the right shows’ quantification of the αSMA^+^ area index for the indicated genotypes. Scale bar indicates 100 μm. D) Panel on right shows Western blot bands for αSMA (green) and GAPDH (red) from lysates generated from indicated genotypes. Histogram on the right shows quantification analysis for Western blots performed on three biological replicates for each genotype. E) Histogram showing qRT-PCR relative expression analysis for transcripts of the indicated genes in the four genotypes used in the study. *Ppia* was used as the normalising gene. In dataplots shown in A), B) and C), square symbols identify males, circle symbols identify females.

**Figure 7:**
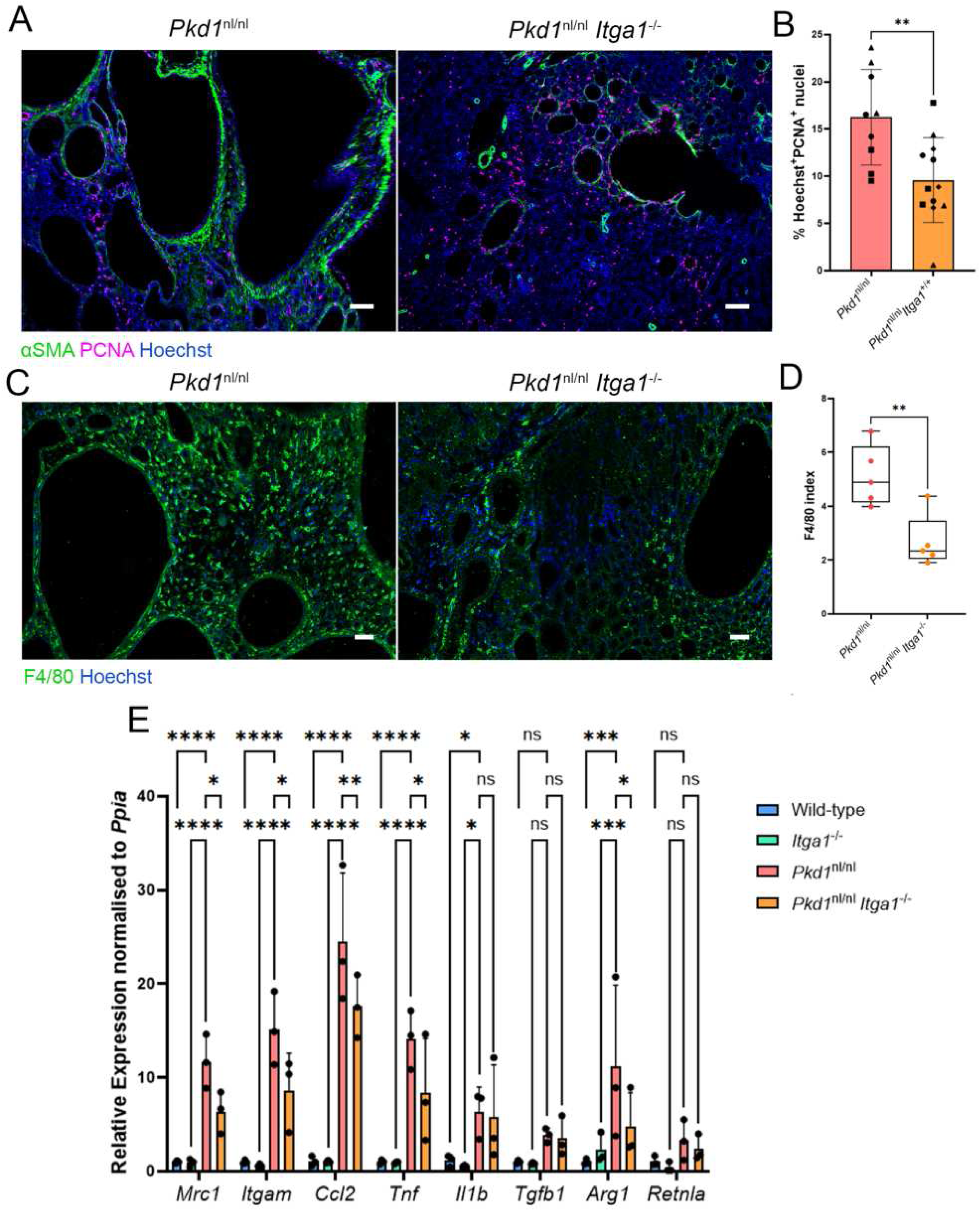
Integrin α1 depletion reduced myofibroblast proliferation and numbers of macrophage in Pkd1^nl/nl^ kidneys. A) Panels show triple staining for αSMA (green), PCNA (magenta) and Hoechst (blue) in *Pkd1*^nl/nl^ (left panel) and *Pkd1*^nl/nl^*Itga1*^−/−^ (right panel) kidneys. Scale bar indicates 100 μm. B) Histogram comparing the % of Hoechst^+^PCNA^+^ nuclei in αSMA^+^ regions only of *Pkd1*^nl/nl^ and *Pkd1*^nl/nl^*Itga1*^−/−^ kidneys. Each symbol represents a biological replicate, for each replicate three random snapshots were taken. C) Panels show staining for F4/80 (green) and Hoechst (blue) in *Pkd1*^nl/nl^ (left panel) and *Pkd1*^nl/nl^*Itga1*^−/−^ (right panel) kidneys. Scale bar indicates 100 μm. D) Box and whisker plot showing the F4/80 index in *Pkd1*^nl/nl^ and *Pkd1*^nl/nl^*Itga1*^−/−^ kidneys. E) Histogram showing qRT-PCR relative expression analysis for transcripts of the macrophage markers and inflammatory genes in the four genotypes used in the study. *Ppia* was used as the normalising gene.

### Study approval

Human tissue used for immunostaining in Figure 3C and S4 were obtained from donations to the PKD Bioresource bank, supported by the UK PKD charity. The Research Ethics Committee code for the tissue bank is 20772. For the experiments using mice, the approaches used were approved by a local AWERB committee and performed under a project licence granted to Richard Naylor by the UK Home Office (PP7091544).

### Data availability

The MS proteomics data will be deposited to the ProteomeXchange Consortium via the PRIDE partner repository[37] upon publication.

## Results

### Cysts in Pkd1^nl/nl^ mice develop fibrosis by P28

To assess the fibrotic phenotype in *Pkd1*^nl/nl^ mice we visualised ECM and myofibroblast markers in 1-day-old (P1) and 4-week-old (P28) mice. These time-points define early (P1) and advanced (P28) stages of cystogenesis in this model[38]. At P1, we observed no difference in kidney weight or body weight between wild-type and *Pkd1*^nl/nl^ mice, but by P28 there was clear separation in these metrics (**Figure S1**). At P1, picrosirius red staining revealed similar collagen staining between wild-type and *Pkd1*^nl/nl^ kidneys, as well as small cysts in *Pkd1*^nl/nl^ (**Figure 1A and 1C**). At P28, we observed large cysts, interstitial expansion and fibrosis as well as pericystic ECM thickening in *Pkd1*^nl/nl^ kidneys (**Figure 1B and 1C**). Myofibroblasts are a key fibrotic cell type in kidney disease, and they express high levels of alpha smooth muscle actin (αSMA). Interestingly, P1 control and *Pkd1*^nl/nl^ kidneys had high levels of αSMA expression in the medullary interstitial region of the kidney, and myofibroblasts were also observed surrounding developing nephrons in the cortex, highlighting a possible role for these cells in nephrogenesis (**Figure 1D and 1F**). At P28, αSMA was restricted to vascular smooth muscle cells enveloping blood vessels in wild-type kidneys (**Figure 1E**), constituting a very small percentage of total kidney area (**Figure 1F**). In *Pkd1*^nl/nl^ kidneys, αSMA was maintained at high levels in the expanded interstitium (**Figure 1E and 1F**). Thus, *Pkd1*^nl/nl^ mice at P1 have no fibrotic niche, but by P28 fibrosis is present.

### MS-based proteomics analysis reveals novel fibrotic features in P28 **Pkd1^nl/nl^** mice

To interrogate global changes in protein abundance between control and *Pkd1*^nl/nl^ kidneys, we performed mass spectrometry-based proteomics. Tryptic peptide preparations were derived from *Pkd1*^nl/nl^ and *Pkd1*^+/+^ littermates at P1 and P28 (**Figure 2A**). Fractionation of lysates into a soluble fraction (SF) and less soluble fraction (LSF) was performed (**see Methods section**), whereby ECM proteins are enriched in the LSF. More than 1800 proteins were detected in each of the samples collected (**Figure 2B**) and principal component analysis revealed clear separation of biological replicates at P28, with less separation at P1 (**Figure 2C**). The three most up- and down-regulated proteins detected in each fraction at P28 were validated by qRT-PCR (**Figure S2A**). The most statistically significant upregulated protein in the P28 SF was plectin, and immunostaining for plectin in P28 kidney sections confirmed its increased expression in pericystic, epithelial and interstitial regions of polycystic kidneys (**Figure S2B**). To investigate pathway enrichment, we generated Voronoi plots using the PantherDB[39] (**Figure 2D and S3**). This analysis showed the top 100 proteins upregulated in our dataset were enriched for RNA metabolism, cytoskeletal and RNA splicing proteins at both P1 and P28. At P28, cell adhesion and extracellular matrix proteins were also enriched. Using the Matrisome database[40], we focussed our analysis on ECM proteins. As expected from results shown in Figure 1, there were only seven SF ECM proteins and four LSF ECM proteins that were differentially expressed at P1, with fibrinogen the main protein group altered (**Figure 2E**). At P28, many detected ECM proteins were differentially expressed. In the soluble fraction, analysis showed 36 upregulated ECM proteins and 11 downregulated ECM proteins. The downregulated ECM proteins included the metalloproteases Meprin1a and 1b, the glomerular basement membrane component Col4a3, and the transglutaminase Tgm1. Upregulated proteins included several proteins previously implicated in fibrogenesis and cystogenesis, including Lcn2, Postn, Lgals3, Tnc, Egf, and Fn1. In the less soluble fraction, four proteins were downregulated, and 49 were upregulated, including many ECM components and pro-fibrotic proteins, such as Lgals3, Lamb1, Col12a1, Hspg2, Lamc1, Sparc, and Fn1. Evidence from Voronoi plots and other gene ontology (GO) analysis suggested a dramatic change in cell-matrix adhesion. Filtering for focal adhesion proteins confirmed a significant change in cell-matrix adhesion (**Figure 2F**). In conclusion, these results show the presence of a fibrotic protein signature at P28 and identify significant global changes in the kidney adhesome of *Pkd1*^nl/nl^ mice.

### Integrin α1 is a marker of kidney fibroblasts and myofibroblasts

Integrins are the principal receptors used by cells to bind to the ECM. We detected four alpha subunits (Itga1, Itga3, Itgam, Itgav) and four beta subunits (Itgb1, Itgb2, Itgb4, Itgb5) in the proteomic analysis, and all were significantly upregulated at P28 (**Figure 3A**). We focused on the integrin α1 subunit as it has not been studied in the context of ADPKD and treatments targeting this subunit are likely to be a well-tolerated due to a lack of any phenotypes associated with its depletion in humans and mice[15]. We reanalysed a single-cell RNA-seq (scRNA-seq) dataset that was performed to investigate the cellular origin of myofibroblasts in human kidney fibrosis[10], and we found that *ITGA1* is enriched in kidney fibroblasts and myofibroblasts, as well as Schwann cells and, at lower levels, the blood vasculature (**Figure 3B and S4**). We also investigated the expression levels of the other collagen-binding integrin subunits (*ITGA2*, *ITGA10* and *ITGA11)* in this dataset. *ITGA2* was expressed in distal tubule and duct segments of the nephron and Schwann cells but was not expressed in mesenchymal cells (**Figure 3B and S4**). *ITGA10* was not expressed in any kidney cell type, whereas *ITGA11* was expressed in myofibroblasts at a lower level than *ITGA1* (**Figure 3B and S4**). To investigate the localisation pattern of integrin α1 in human ADPKD tissue, we obtained control normal healthy kidney tissue, early predialysis-stage ADPKD kidney tissue (E-ADPKD) and end-stage (in kidney failure) ADPKD kidney tissue (ES-ADPKD) from the PKD Charity Bioresource bank. Co-immunofluorescence for integrin α1 and αSMA showed integrin α1 labelled mesenchymal cells in the interstitium of normal healthy kidney tissue and co-labels αSMA^+^ cells in E-ADPKD tissue (**Figure 3C and S5**). Quantification of Integrin α1^+^ and αSMA^+^ area showed an increase in E-ADPKD and ES-ADPKD tissue (**Figure 3D**). We suggest this is primarily due to interstitial expansion as fibrotic scarring worsens with disease progression (**Figure S5**). We also analysed the large kidney scRNA-seq dataset generated by the Kidney Precision Medicine Project[41], which showed *ITGA1* expression in mesenchymal and endothelial cell groups, as well as parietal epithelial cells (**Figure 3E**). This analysis revealed *ITGA1* is upregulated in fibroblasts and myofibroblasts in acute kidney injury (AKI, n=14) and chronic kidney disease (CKD, n= 37) in comparison to healthy reference (n=26) (**Figure 3E**). *ITGA11* also showed an upregulation in myofibroblasts in AKI and CKD, but to a lesser degree than *ITGA1* (**Figure S6**). Thus, integrin α1 is a marker of mesenchymal cells in the kidney interstitium, most likely resident fibroblasts in normal healthy kidney and activated myofibroblasts in diseased kidneys.

### Integrin α1 depletion reduced pathologies associated with **Pkd1^nl/nl^** mice

Given integrin α1 is highly expressed in kidney interstitial mesenchymal cells and is upregulated in *Pkd1*^nl/nl^ mice, we hypothesised that integrin α1 has a role in the pathogenesis of ADPKD. To test this, we crossed global *Itga1*^−/−^ mice with the *Pkd1*^nl/nl^ model of ADPKD (**Figure 4A**). Kidneys isolated from *Pkd1*^nl/nl^*Itga1*^−/−^ mice were much smaller than those isolated from *Pkd1*^nl/nl^ mice but had a cystic appearance and were larger than wild-type and *Itga1*^−/−^ control kidneys (**Figure 4B**). Analysis of the 2-kidney weight to body weight ratio (2KW:BW) showed *Pkd1*^nl/nl^*Itga1*^−/−^ kidneys (n=7, _x_LJ = 4.83%±1.05) had a 48.9% reduction in 2KW:BW compared to *Pkd1*^nl/nl^ kidneys (n=11, _x_LJ = 9.45%±1.92) (**Figure 4C**). However, their body weights were similar to *Pkd1*^nl/nl^ mice, which was low compared to wild-type and *Itga1*^−/−^ mice (**Figure 4D**). Furthermore, *Pkd1*^nl/nl^*Itga1*^−/−^ 2KW:BW was 3.6x greater than *Itga1*^−/−^ mice (n=8, _x_LJ =1.36%±0.21) and 3.5x greater than wild-type mice (n=10, _x_LJ = 1.38%±0.16). Kidney dysfunction was measured by determining blood urea nitrogen (BUN) levels and the urinary albumin creatinine ratio (uACR). *Pkd1*^nl/nl^*Itga1*^−/−^ mice had a 21.03% decline in BUN and a 39.45% decline in uACR compared to *Pkd1*^nl/nl^ mice (n=8, **Figure 4E and 4F**). BUN levels in *Pkd1*^nl/nl^*Itga1*^−/−^ mice were still elevated compared to wild-type (+122.44%) and *Itga1*^−/−^ (+108.49%) mice (n=8, **Figure 4E**) and uACR levels in *Pkd1*^nl/nl^*Itga1*^−/−^ mice were increased +47.45% compared to wild-type and +20.96% compared to *Itga1*^−/−^ (n=8, **Figure 4F**). Taken together, these results show that depletion of integrin α1 in *Pkd1*^nl/nl^ mice ameliorates but does not prevent kidney disease in this model of ADPKD. This finding suggest integrin α1 contributes to kidney pathogenesis associated with the *Pkd1*^nl/nl^ genotype.

### Integrin α1 depletion reduces cystogenesis and fibrosis in **Pkd1^nl/nl^** mice

We next performed histological analysis on isolated kidneys to determine the effect of integrin α1 depletion in *Pkd1*^nl/nl^ mice. Cysts were still present in all *Pkd1*^nl/nl^*Itga1*^−/−^ kidneys analysed by hematoxylin and eosin (H&E) staining (**Figure 5A**). However, the cystic index (**Figure 5B**) and average cyst size (**Figure 5C**) were reduced 25.7% and 40.03% in *Pkd1*^nl/nl^*Itga1*^−/−^ kidneys compared to *Pkd1*^nl/nl^ kidneys, respectively. The average number of cysts was not changed in these two groups (**Figure 5D**). The number of glomeruli in single section counts was reduced by 55% (n=6, *p*=0.0007) in *Pkd1*^nl/nl^ kidneys, but was not significantly changed in *Pkd1*^nl/nl^*Itga1*^−/−^ kidneys (n=6, *p*=0.411) compared to control kidneys (**Figure 5E**). Closer observation of *Pkd1*^nl/nl^*Itga1*^−/−^ kidneys showed non-cystic kidney parenchyma was well maintained compared to *Pkd1*^nl/nl^ kidneys, with reduced interstitial expansion (**Figure 5F**). Taken together, these findings demonstrate loss of integrin α1 reduced tubular atrophy and scarring in *Pkd1*^nl/nl^ kidneys.

These histological observations combined with our expression data showing integrin α1 labels kidney interstitial mesenchymal cells, suggested integrin α1 might be involved in fibrogenesis in *Pkd1*^nl/nl^ kidneys. To further investigate the impact of integrin α1 depletion on interstitial fibrosis in *Pkd1*^nl/nl^ mice, we stained kidney slides with picrosirius red (PSR) staining. Deconvolution of brightfield PSR images to specifically measure collagen labelling showed a reduction in interstitial ECM deposition in *Pkd1*^nl/nl^*Itga1*^−/−^ kidneys compared to *Pkd 1*^nl/nl^ kidneys (**Figure 6A**, for whole kidney views see **Figure S7**). Analysis of PSR red fluorescence identified a similar reduction in collagen deposition in *Pkd1*^nl/nl^ kidneys depleted of integrin α1 (**Figure 6B**, for whole kidney views see **Figure S7**). We noted that pericystic collagen deposition persisted in *Pkd1*^nl/nl^*Itga1*^−/−^ kidneys, suggesting that the reduction in ECM deposition observed in these kidneys was likely due to restricted interstitial fibrosis. Given myofibroblasts are the major pro-fibrotic cell type that deposit ECM in the kidney[10], and we have shown integrin α1 is a marker of myofibroblasts in human ADPKD, we analysed αSMA to see what effect integrin α1 depletion had on this cell type. We found that the total area of αSMA expression was reduced in *Pkd1*^nl/nl^*Itga1*^−/−^ kidneys (n=8) compared to *Pkd1*^nl/nl^ kidneys (n=8, **Figure 6C**). αSMA^+^ staining was observed in the mesenchyme of *Pkd1*^nl/nl^*Itga1*^−/−^ kidneys, which was not observed in wild-type (n=6) and *Itga1*^−/−^ (n=6) control kidneys (**Figure 6C**, for whole kidney views see **Figure S8**). This result suggests fibroblasts can transdifferentiate to myofibroblasts in *Pkd1*^nl/nl^*Itga1*^−/−^ kidneys, but there are fewer of these cells. Western blot analysis of kidney lysates also showed αSMA levels in *Pkd1*^nl/nl^*Itga1*^−/−^ kidneys were reduced compared to *Pkd1*^nl/nl^ kidneys but increased when compared to wild-type and *Itga1*^−/−^ control kidneys (**Figure 6D**). qRT-PCR analysis for fibrotic markers *Acta2*, *Col1a1*, *Fn1* and *SerpinE1* showed these markers were reduced in *Pkd1*^nl/nl^*Itga1*^−/−^ kidneys compared to *Pkd1*^nl/nl^ kidneys, but still increased compared to wild-type and *Itga1*^−/−^ controls (**Figure 6E**). Given integrin α1 depletion reduces cell proliferation in dermal fibroblasts [22], we speculated loss of integrin α1 may have reduced or prevented expansion of the kidney myofibroblast pool in *Pkd1*^nl/nl^ kidneys. We performed co-labelling of αSMA and the cell proliferation marker PCNA to measure the number of myofibroblasts that were dividing in the mesenchyme (**Figure 7A and S9**). We observed a 40.95% reduction in the proportion of co-labelled PCNA^+^/αSMA^+^ cells in *Pkd1*^nl/nl^*Itga1*^−/−^ kidneys (9.59%±4.49, n=4) compared to *Pkd1*^nl/nl^ kidneys (16.25%±5.06, n=3, **Figure 7B**).

Taken together, these results suggest that fibroblast-to-myofibroblast transdifferentiation is possible in *Pkd1*^nl/nl^ kidneys depleted of integrin α1, but these cells are fewer in number (most likely due to reduced proliferation) and have lowered expression for pro-fibrotic markers.

### Integrin **α**1 depletion reduced the number of macrophage in the **Pkd1^nl/nl^** kidney

An important component of the chronic response to cyst growth in ADPKD is the involvement of invasive and tissue-resident leukocytes. Our proteomics analysis showed increased expression of the macrophage integrin αM subunit (**Figure 3A**). In addition to integrin αM, Integrin α1 has also been reported to be expressed on circulating monocytes and activated macrophages[27, 42], although our scRNA-seq analyses do not corroborate this (**Figures 3B and 3E**). Regardless, macrophages are the major phagocytic cell type in the body and are closely associated with myofibroblasts in many fibrotic disease contexts[9, 43]. To observe the effect of integrin α1 depletion on the number of leukocytes in *Pkd1*^nl/nl^ kidneys, we performed immunofluorescence with the F4/80 macrophage marker. We observed a reduction in the number of F4/80^+^ macrophage in *Pkd1*^nl/nl^*Itga1*^−/−^ kidneys compared to *Pkd 1*^nl/nl^ kidneys (**Figure 7C and 7D**). qRT-PCR analysis also showed a reduction in the macrophage markers *Mrc1* and *Itgam* in *Pkd1*^nl/nl^*Itga1*^−/−^ kidneys, as well as the *Ccl2* chemokine secreted by fibroblasts to attract macrophages (**Figure 7E**). Reduced expression of inflammatory markers *Tnf* and *Arg1* was also detected after integrin α1 depletion, but *Il1b, Tgfb1,* and *Retnla* expression remained unchanged (**Figure 7E**).

### Integrin α1 depletion reduced primary fibroblast pro-fibrogenic phenotype

To further investigate the role of integrin α1 depletion in the fibroblast response to ADPKD, we isolated primary renal fibroblasts from *Pkd1*^nl/nl^ and *Pkd1*^nl/nl^*Itga1*^−/−^ mice. With immunofluorescence we found 92.96%±2.79% of primary fibroblasts isolated from *Pkd1*^nl/nl^ kidneys were αSMA^+^ (n=4, **Figure 8A and 8B**). In *Pkd1*^nl/nl^*Itga1*^−/−^ kidneys, 80.03%±3.86% of primary fibroblasts were αSMA^+^ (n=3, **Figure 8A and 8B**). These results indicate a statistically significant reduction in the number of αSMA^+^ primary fibroblasts from *Pkd1*^nl/nl^*Itga1*^−/−^ kidneys (*p*=0.0035). No overall significant differences were observed in proliferation rates between primary fibroblasts isolated from *Pkd1*^nl/nl^ and *Pkd1*^nl/nl^*Itga1*^−/−^ fibroblasts (**Figure 8C**). This contrasts with PCNA^+^ observations *in vivo*, suggesting phenotypic change when these cells are cultured *in vitro*. With contractility assays we found *Pkd1*^nl/nl^ primary fibroblasts reduced the size of collagen hydrogels by 46.31%±7.1 compared to no cell controls, whereas *Pkd1*^nl/nl^*Itga1*^−/−^ primary fibroblasts collagen hydrogels contracted by 5.86%±7.47 (**Figure 8D**). qRT-PCR analysis of *Pkd1*^nl/nl^ and *Pkd1*^nl/nl^*Itga1*^−/−^ primary fibroblasts showed fibrotic markers *Col1a1*, *Fn1* and *SerpinE1* were reduced by integrin α1 depletion (**Figure 8E**). Taken together, these results support *in vivo* findings that pro-fibrogenic actions of myofibroblasts in ADPKD kidneys are augmented by integrin α1.

**Figure 8:**
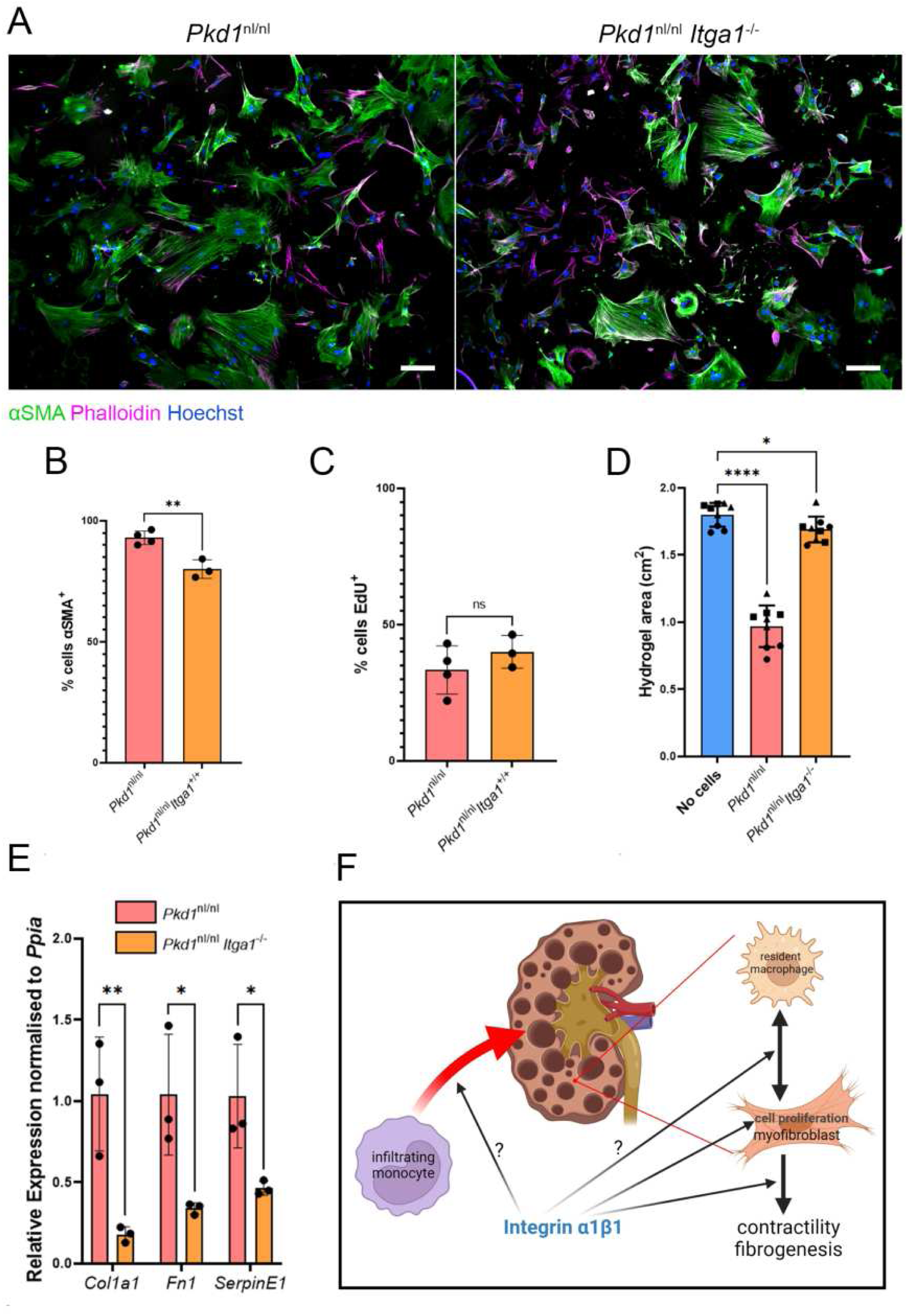
Profibrogenic markers were reduced in primary fibroblasts cultures taken from. Pkd1^nl/nl^Itga1^−/−^ **kidneys** A) Panels show triple staining for αSMA (green), phalloidin (magenta) and Hoechst (blue) in primary fibroblast cultures taken from *Pkd1*^nl/nl^ (left panel) and *Pkd1*^nl/nl^*Itga1*^−/−^ (right panel) kidneys. Scale bar indicates 100 μm. B) and C) Histograms indicating the % of αSMA^+^ cells (B) and % EdU^+^ nuclei in primary fibroblast cultures taken from *Pkd1*^nl/nl^ and *Pkd1*^nl/nl^*Itga1*^−/−^ kidneys. D) Histogram showing the contractility of collagen hydrogels seeded with no cells, *Pkd1*^nl/nl^ primary fibroblasts and *Pkd1*^nl/nl^*Itga1*^−/−^ primary fibroblasts. E) Histogram showing qRT-PCR relative expression analysis for transcripts of the fibrotic markers in *Pkd1*^nl/nl^ versus *Pkd1*^nl/nl^*Itga1*^−/−^ primary fibroblasts. *Ppia* was used as the normalising gene. F) Schematic model for the proposed mechanisms by which integrin α1 promotes kidney fibrosis.

## Discussion

The goal of this study was to utilise our expertise in MS-based proteomics to analyse *Pkd1*^nl/nl^ mice and detect hitherto uncharacterised molecular changes in ADPKD. This was especially important as most analyses in ADPKD research have relied upon transcriptomic approaches. We identified a general increase in cell adhesion proteins, including multiple integrin receptors, in our proteomics dataset. Previous studies have investigated the roles of the integrin β1 subunit[44, 45] and integrin-linked kinase[46] signalling in cyst growth and fibrosis in ADPKD. Roles for integrins α_v_β3[47], α_v_β5[48], α_v_β6[49], α3β1[50, 51], α4β1/4[52, 53] and α_m_β2[54] in fibrosis have also been studied. A general theme from these studies is that integrins augment fibrosis by promoting both the immune response to tissue-specific inflammation and the fibrogenic actions of myofibroblasts. We detected all these integrin subunits with increased abundance in the *Pkd1*^nl/nl^ proteomics in addition to increased integrin α1, which binds integrin β1 to form the heterodimeric integrin α1β1 receptor. Given the lack of any obvious phenotypes associated with healthy and/or injured *Itga1* null mice[15], it is likely that any therapy targeting integrin α1β1 will be well tolerated. Therefore, we wished to investigate if integrin α1 has a functional role in ADPKD pathogenesis. Analysis of *Pkd1*^nl/nl^*Itga1*^−/−^ mice showed a dramatic reduction in kidney size and kidney dysfunction, identifying integrin α1 as a mediator of ADPKD pathogenesis. Strikingly, we observed a significant reduction in interstitial fibrosis in *Pkd1*^nl/nl^*Itga1*^−/−^ kidneys. The non-cystic parenchyma of these mice was relatively well maintained; the number of glomeruli was similar to wild-type controls and interstitial expansion was relatively restricted. We believe this reduced fibrotic response is the causal factor that improved kidney function in *Pkd1*^nl/nl^ kidneys depleted of integrin α1.

Using immunofluorescence and scRNA-seq data[10], we showed integrin α1 is expressed and localised in mesenchymal cells that populate the kidney interstitium. Previously, integrin α1 expression in mice has been reported in the glomerular mesangium, in embryonic stage proximal tubules and in the kidney interstitium[27, 55] using a specific antibody[56]. In *Pkd1*^nl/nl^*Itga1*^−/−^ kidney primary fibroblast cultures, integrin α1 depletion reduced myofibroblast contractility and expression of pro-fibrotic genes (*Col1a1*, *Fn1* and *SerpinE1*). Given the expression profile of integrin α1 in fibroblasts and myofibroblasts and the reduced fibrotic phenotype upon its depletion, it is likely that integrin α1 is directly involved in the tissue repair response performed by myofibroblasts. Intriguingly, we only observed a marginal reduction in αSMA immunofluorescence in primary cultured fibroblasts taken from *Pkd1*^nl/nl^ and *Pkd1*^nl/nl^*Itga1*^−/−^ kidneys. It is likely that a 2D *in vitro* environment is sufficient to induce a myofibroblast phenotype, which might explain high levels of αSMA expression in both samples. Despite this, integrin α1 depletion reduced contractility in primary kidney fibroblasts of our ADPKD mouse model. Similar observations have been observed in activated liver stellate cells (a type of myofibroblast), where anti-integrin α1 blocked contraction by 70%[56]. Integrin α1 expression is present in normal stellate cells and persists when they are activated in response to injury[56], a finding that correlates with our observations in human healthy and ADPKD kidney tissue. Overexpression of integrin α1 in cultured rat glomerular mesangial cells has also been shown to increase contractility of these cells[57]. In heart myofibroblasts, contractility strength is not dependent on the binary presence or absence of αSMA; as fibrosis progresses, so too does contractility strength[58]. Focal adhesions are integral to αSMA-mediated contractions, and increased contractility further enhances ECM deposition and fibrogenesis in a positive feedback loop[59]. Therefore, our finding that integrin α1-depleted primary myofibroblasts express αSMA but are less contractile and less fibrogenic might imply this positive feedback loop is minimised or blocked.

Our results contrast with previous observations showing a lack of integrin α1 leads to increased fibrosis in the kidney and skin. In dermal skin wounding, mice lacking integrin α1 have fewer fibroblasts due to lowered rates of proliferation[22], but overall more fibrosis[21]. And in models of glomerular injury, *Itga1*-null mice have increased sclerosis[18, 19]. One possible explanation for this discrepancy is that the glomerular injury induced by adriamycin or streptozotocin primarily affects mesangial cells within the glomerulus of the kidney. In this setting, integrin α1 appears to be protective against excessive ECM deposition. Glomerular injury is usually not associated with ADPKD, and nephrotic range proteinuria is unusual in patients with ADPKD. Thus, the role of integrin α1β1 in tubulopathy (such as ADPKD) is distinct to that exerted in glomerulopathy. However, interstitial fibrosis is exacerbated in *Itga1*^−/−^ mice in the UUO model of fibrosis[20]. Fibrosis develops in the UUO model within days, which is more reminiscent of an acute kidney injury rather than chronic kidney disease. Therefore, it is possible that in a more clinically relevant setting, where interstitial fibrosis develops over a longer period because of chronic kidney disease, integrin α1β1 inhibition may be beneficial.

Our primary fibroblast culture experiments highlighted that integrin α1-depletion is unable to prevent fibroblast-to-myofibroblast transdifferentiation (based on αSMA expression). This is consistent with our *in vivo* findings where much of the tubulointerstitium of *Pkd1*^nl/nl^*Itga1*^−/−^ kidneys was weakly αSMA^+^. As discussed above, the reduced contractility and expression of pro-fibrotic markers in integrin α1 depleted myofibroblasts is likely to be a significant factor for reduced fibrosis in *Pkd1*^nl/nl^*Itga1*^−/−^ kidneys. However, additional factors may also be involved, including the role of integrin α1β1 on leukocytes that invade the inflamed kidney and promote fibrosis. We observed a 50% reduction in F4/80^+^ leukocytes in *Pkd1*^nl/nl^*Itga1*^−/−^ kidneys. These cells are most likely macrophages, the origins of which are either resident embryonic macrophages or infiltrating monocytes. Integrin α1β1 has been reported to be expressed on activated macrophages[42] in the kidney[27]. However, in the scRNA-seq analyses used in this study, the only immune cell type expressing *ITGA1* in the kidney were T cells (**Figures 3B and 3E**), thus further study is required to clarify this discrepancy. Inhibition of integrin α1 on immune cells is recognised to reduce their ability to respond to inflammatory signals and anti-integrin α1 treatments have been used to dampen inflammation in diseases such as rheumatoid arthritis and colitis[60, 61]. These treatments also reduce glomerular and tubulointerstitial scarring in a rat model of glomerulonephritis[62]. In a peripheral lesion murine model of acute inflammation, integrin α1β1 promoted retention of macrophages at the site of injury[63]. Integrin α1 has also been shown to promote monocyte infiltration into the fibrotic kidney by binding to collagen XIII on inflamed vascular endothelium[64]. As well as a reduction in macrophage numbers in *Pkd1*^nl/nl^*Itga1*^−/−^ kidneys, we also observed a different spatial organisation of these cells. In *Pkd1*^nl/nl^ kidneys, macrophages were observed throughout the expanded interstitium, whereas in *Pkd1*^nl/nl^*Itga1*^−/−^ kidneys macrophages were restricted to pericystic regions that are likely to be the primary injury sites. Macrophages are important mediators of fibrosis through proximal and prolonged interaction with myofibroblasts[65]. These cells are thought to create a two-cell circuit, where close and extended interaction ensures myofibroblasts become fully activated[43]. There is also some genetic fate-mapping evidence that macrophages can transdifferentiate to myofibroblasts[66]. Therefore, future investigations into how integrin α1β1 affects fibrosis in chronic tubulopathies might focus on its role in the (myo)fibroblast-to-macrophage circuit.

Cyst expansion in *Pkd1*^nl/nl^ kidneys was reduced after integrin α1 depletion. Given *Itga1* is not expressed in expanding cyst epithelium, we suggest the reduced kidney fibrosis in these mice is the cause of precluded cyst expansion. Recently, it was shown that depleting myofibroblasts in the *Pkd1*^RC/RC^ murine model of ADPKD reduced kidney cyst growth[7]. The mechanism by which myofibroblasts affect cyst growth is likely to be multifaceted. Myofibroblasts may promote epithelial cell proliferation by regulating ECM deposition and turnover, and growth factor production. It is possible that the contractility of myofibroblasts induces stretch-activated proliferation in epithelial cells lining cysts. We note that, even though interstitial fibrosis was perturbed in *Pkd1*^nl/nl^*Itga1*^−/−^ kidneys, the pericystic ECM persisted, suggesting different mechanisms of matrix production between fibroblasts and macrophages that are pericystic versus those that are positioned in the non-cystic kidney parenchyma. Integrin α1β1 has previously been shown to promote cell proliferation in dermal fibroblasts[22] and in a variety of cancers[23, 67] through activation of Ras/ERK signalling. We showed integrin α1 depletion reduced the number of kidney *Pkd1*^nl/nl^ myofibroblasts expressing the proliferation marker PCNA *in vivo*. Measurement of rates of proliferation in primary fibroblasts *in vitro* using EdU labelling showed no significant change, but it is possible that this is due to the artificial cell culture environment. The mouse model of ADPKD used here (*Pkd1*^nl/nl^) yields an early onset of cystogenesis. Given cell proliferation is increased in embryonic stages, and integrin α1β1 is involved in cell proliferation, it is vital that the impact of integrin α1 depletion or integrin α1β1 inhibition in later/slower progressing models of ADPKD is investigated to provide additional evidence that integrin α1β1 is a viable drug target.

Overall, our findings suggest targeting integrin α1β1 could be a viable therapeutic approach to limit interstitial fibrosis in ADPKD. Combining our findings with those in the literature, we propose integrin α1β1 promotes kidney fibrosis by regulating the fibrogenic actions and rates of proliferation of kidney fibroblasts, as well as enabling the immune response to inflammation (**Figure 8F**). Given the lack of phenotypes associated with integrin α1 depletion in healthy mice and the lack of human diseases directly attributed to variants in *ITGA1*, new ADPKD therapies targeting this receptor are likely to be well tolerated. Future studies will test the efficacy of using integrin α1β1 inhibitors to reduce kidney fibrosis and will determine the molecular mechanisms by which integrin α1 promotes fibrogenesis.

## Author contributions

RWN provided the overall study concept, RWN/CG/RL/AP contributed specific experimental design, RWN/CG carried out the experiments, I-HL re-analysed the scRNA-seq data, all authors contributed to analysis of the results and writing of the manuscript.

## Acknowledgements

This work was supported by a Kidney Research UK Intermediate Fellowship (to R.N.), Wellcome Centre for Cell-Matrix Research: 203128/A/16/Z, Wellcome Discovery Research Platform for Cell-Matrix Biology: 226804/Z/22/Z, and, in part, by National Institutes of Health (NIH) grant R01-DK119212 (A.P.); Department of Veterans Affairs Merit Review 1I01BX002025 (A.P.). A.P. is the recipient of a Department of Veterans Affairs Senior Research Career Scientist Award IK6 BX005240. We thank Joseph Stewart and Rachel Mensforth for maintaining the mouse lines used in this study. The Bioimaging Facility microscopes used in this study were purchased with grants from BBSRC, Wellcome and the University of Manchester Strategic Fund. Special thanks go to James Bagnall, Peter March and Steven Marsden for their help with the microscopy.

## Figure legends

Figures S1: A) Box and violin plots showing 2 kidney weight (grams), body weight (grams) and 2 KW:BW ratio of P1 mice used for tissue collection in the proteomics analysis. B) Box and violin plots showing 2 kidney weight (grams), body weight (grams) and 2 KW:BW ratio of P28 mice used for tissue collection in the proteomics analysis.

Figure S2: A) Histogram showing qRT-PCR relative expression analysis for transcripts of the three most upregulated and down-regulated proteins in the proteomics datasets from the SF and LSF used in the study. *Ppia* was used as the normalising gene. B) Panels show immunofluorescence staining for plectin in wild-type (top panels) and *Pkd1*^nl/nl^ (bottom panels) kidneys. Panels on the right are higher magnification confocal images of the indicated genotypes. Scale bars in left panels indicate 500 μm, scale bars in the right panels indicate 50 μm.

Figure S3: Panels show Voronoi plots for Gene names and PantherDB pathway analysis for the SF and LSF comparing Pkd1nl/nl kidneys to wild-type kidneys at P1 and P28. The analysis only includes the top 100 upregulated proteins.

Figure S4: A)-C) Shows uniform manifold approximation and projection distributions of a human kidney scRNA-seq dataset illustrating specific cellular populations (A), ECM and collagen score (B), and specific genes – ITGA1, ITGA2, ITGA10, ITGA11 (C).

Figure S5: A) Panels show αSMA (green) and integrin α1 immunostaining with Hoechst nuclear staining (blue) on human normal healthy kidney tissue (top panels), E-ADPKD kidney tissue (middle panels) or ES-ADPKD kidney tissue (bottom panels). Left panels show tiled confocal large views of human kidneys, scale bar indicates 100 μm. Right panels show super-resolution high magnification views of human kidneys, scale bar indicates 20 μm.

Figure S6: Dot plots of *ITGA1/ITGA2/ITGA10/ITGA11*expression in a scRNA-seq dataset published by the kidney precision medicine project. For cell cluster abbreviation definitions, see Methods section.

Figure S7: Whole kidney views of Picrosirius red (PSR) stained kidneys of the indicated genotypes. Kidneys on the left are imaged under brightfield, kidney on the right side are image in the red fluorescent channel. Scale bars indicate 500 μm.

Figure S8: Whole kidney views of αSMA immuno-stained and Hoechst-stained kidneys of the indicated genotypes.

Figure S9: Schematic describing the FIJI pipeline for determining the number of PCNA^+^ nuclei specifically in the region of αSMA^+^ staining to determine the number of myofibroblasts that have the proliferative signal in kidney samples.

