## Supplementary figures and images for "Integrin alpha1 beta1 promotes interstitial fibrosis in a mouse model of polycystic kidney disease"

### Supplemental Figures

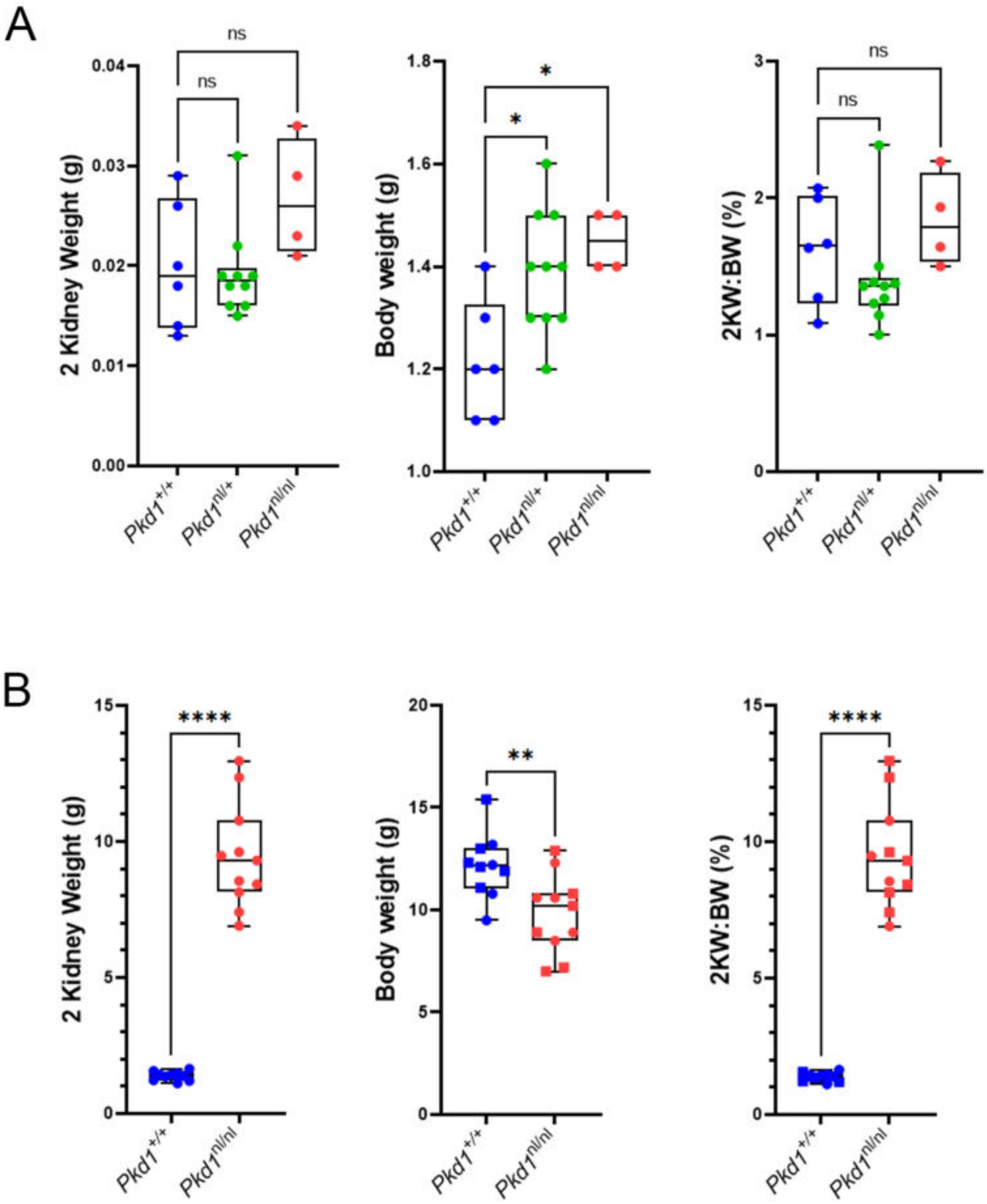

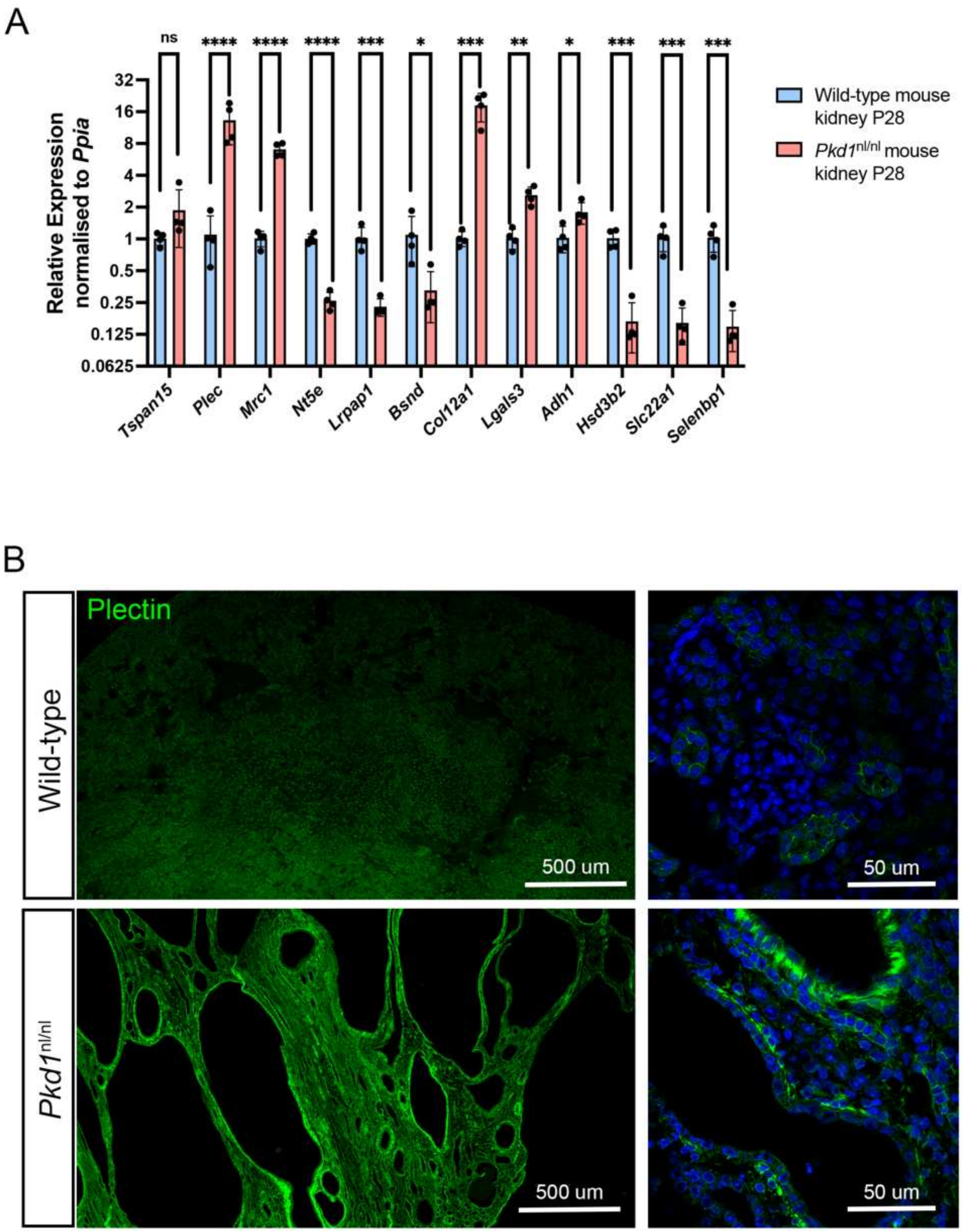

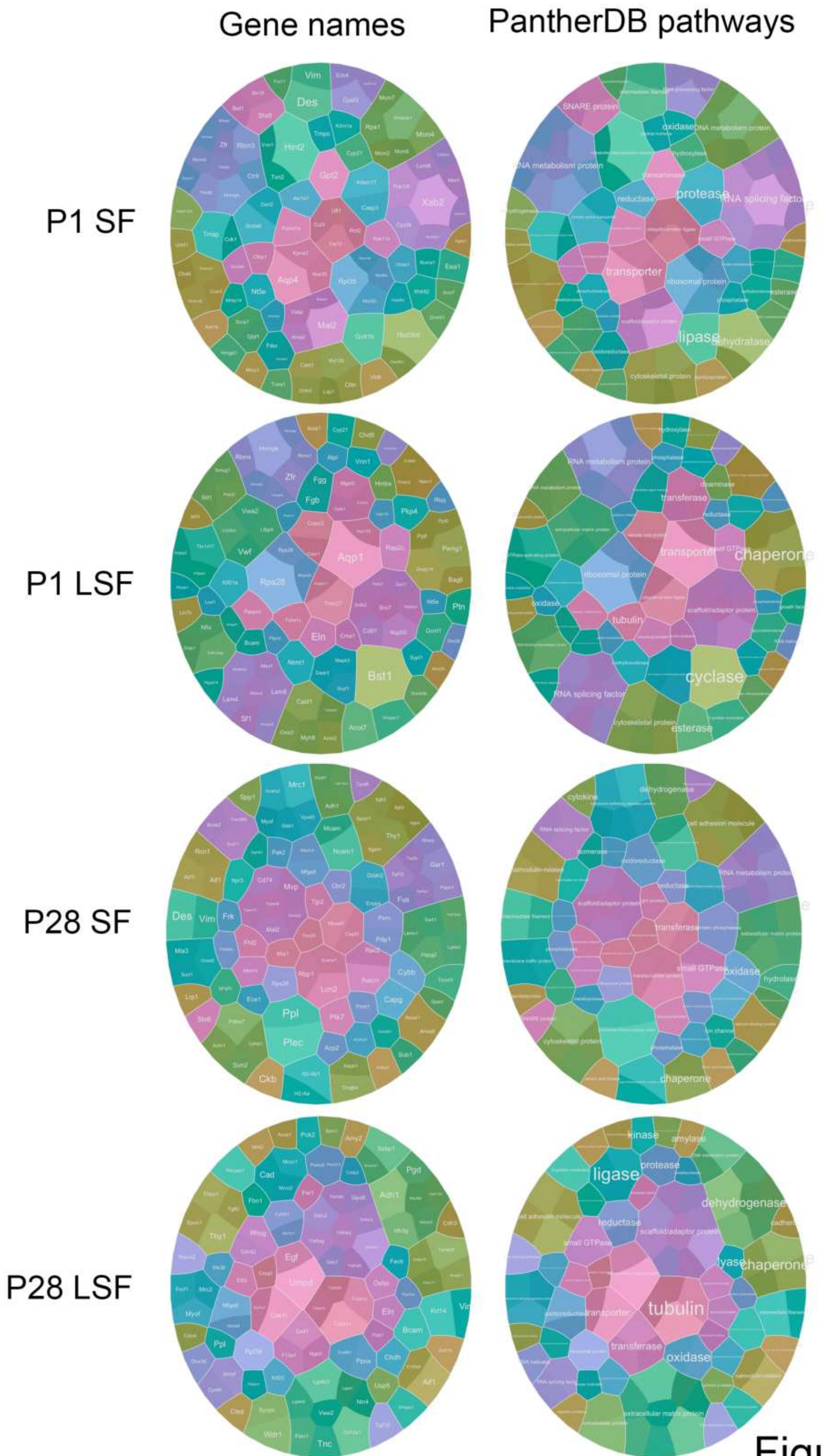


Figure S3

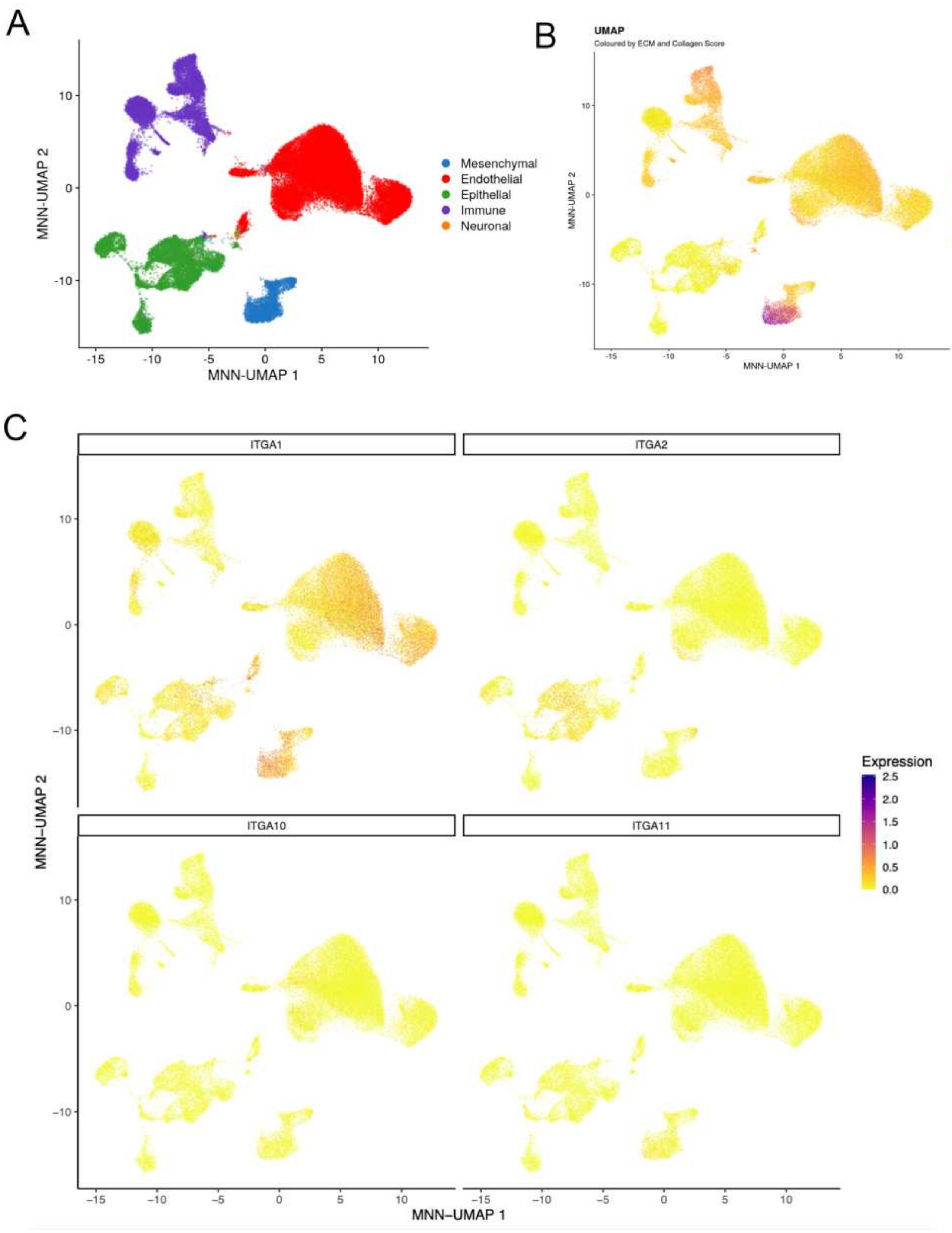

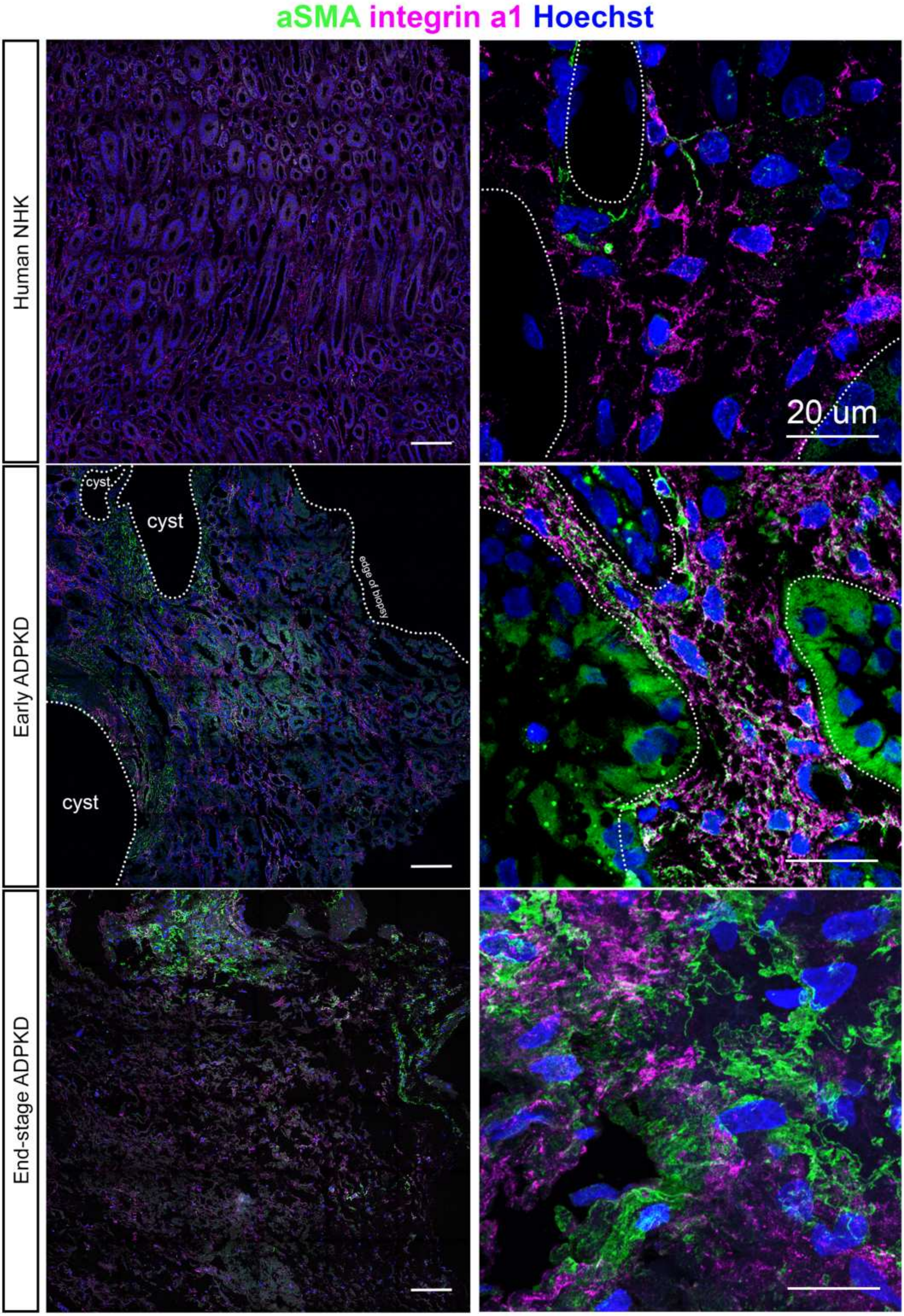

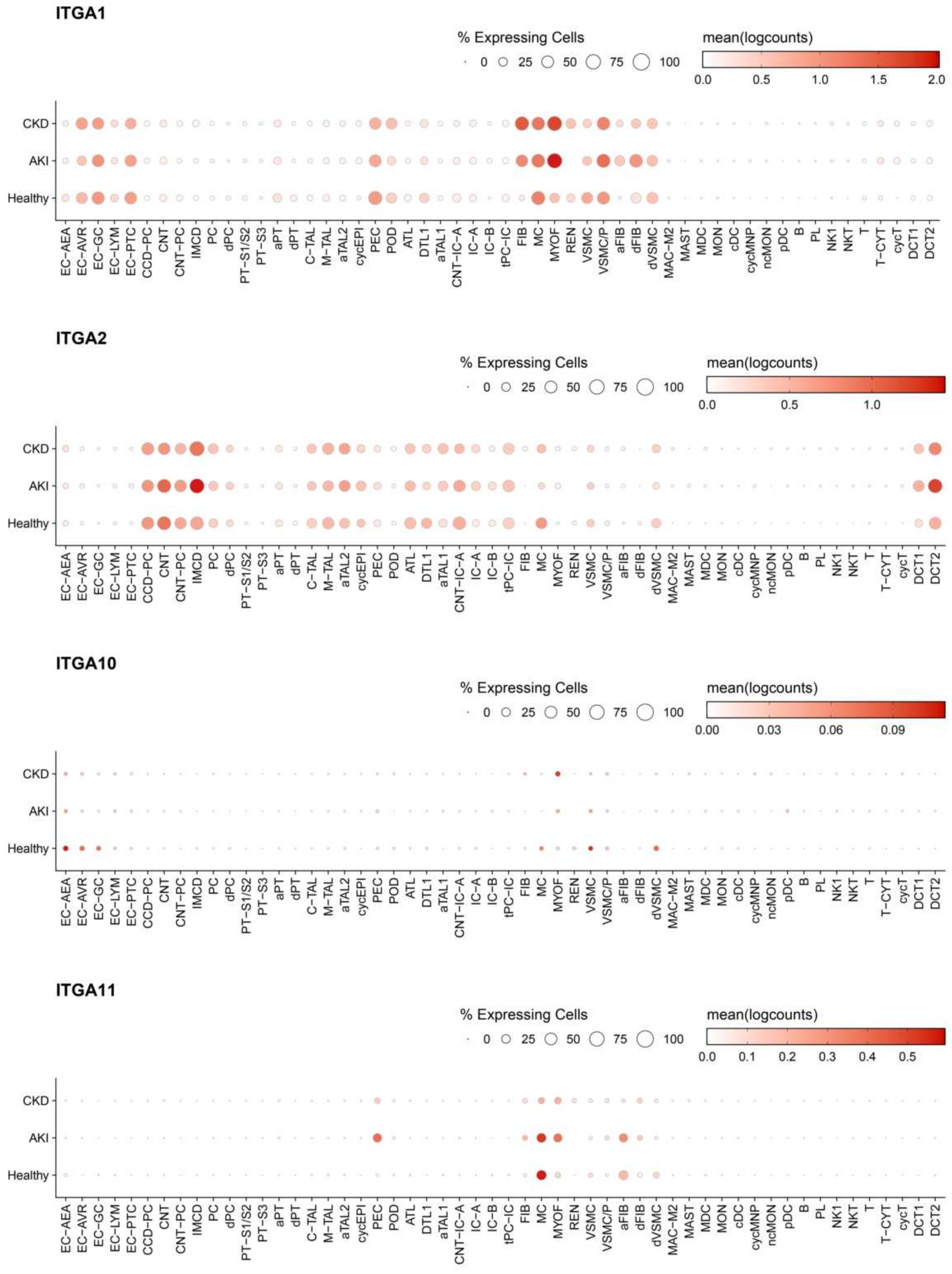

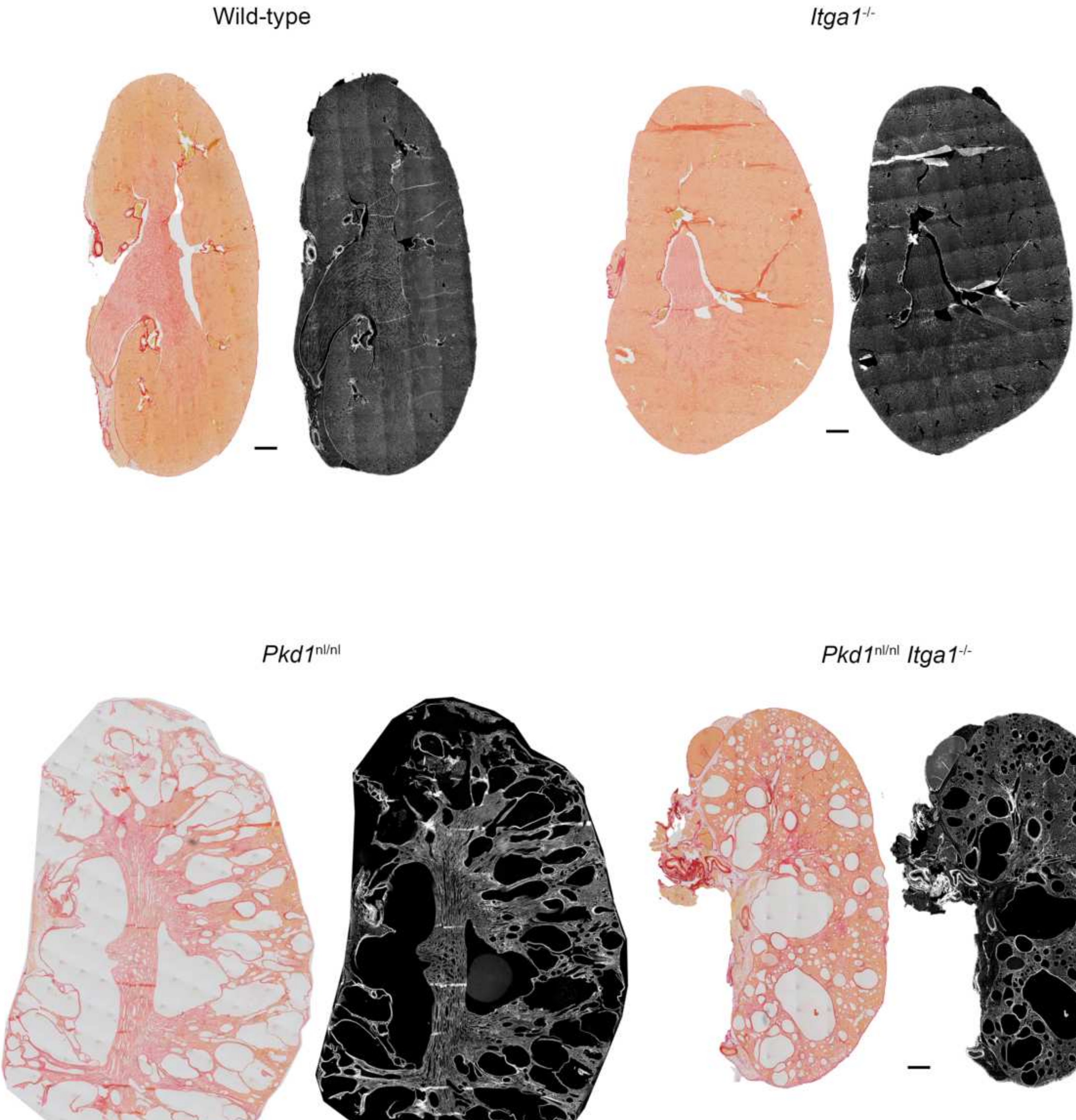


Figure S7

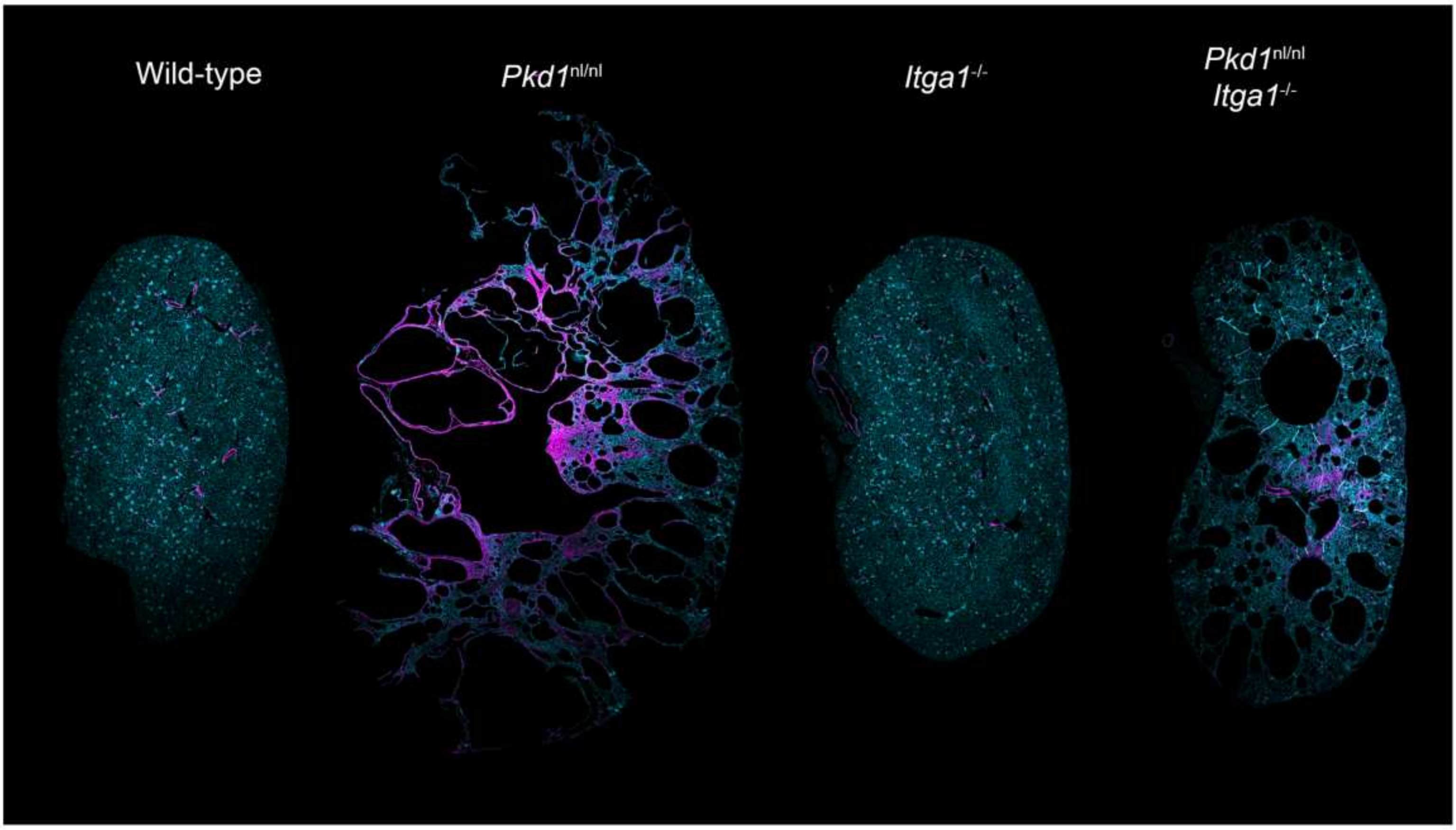


Figure S8

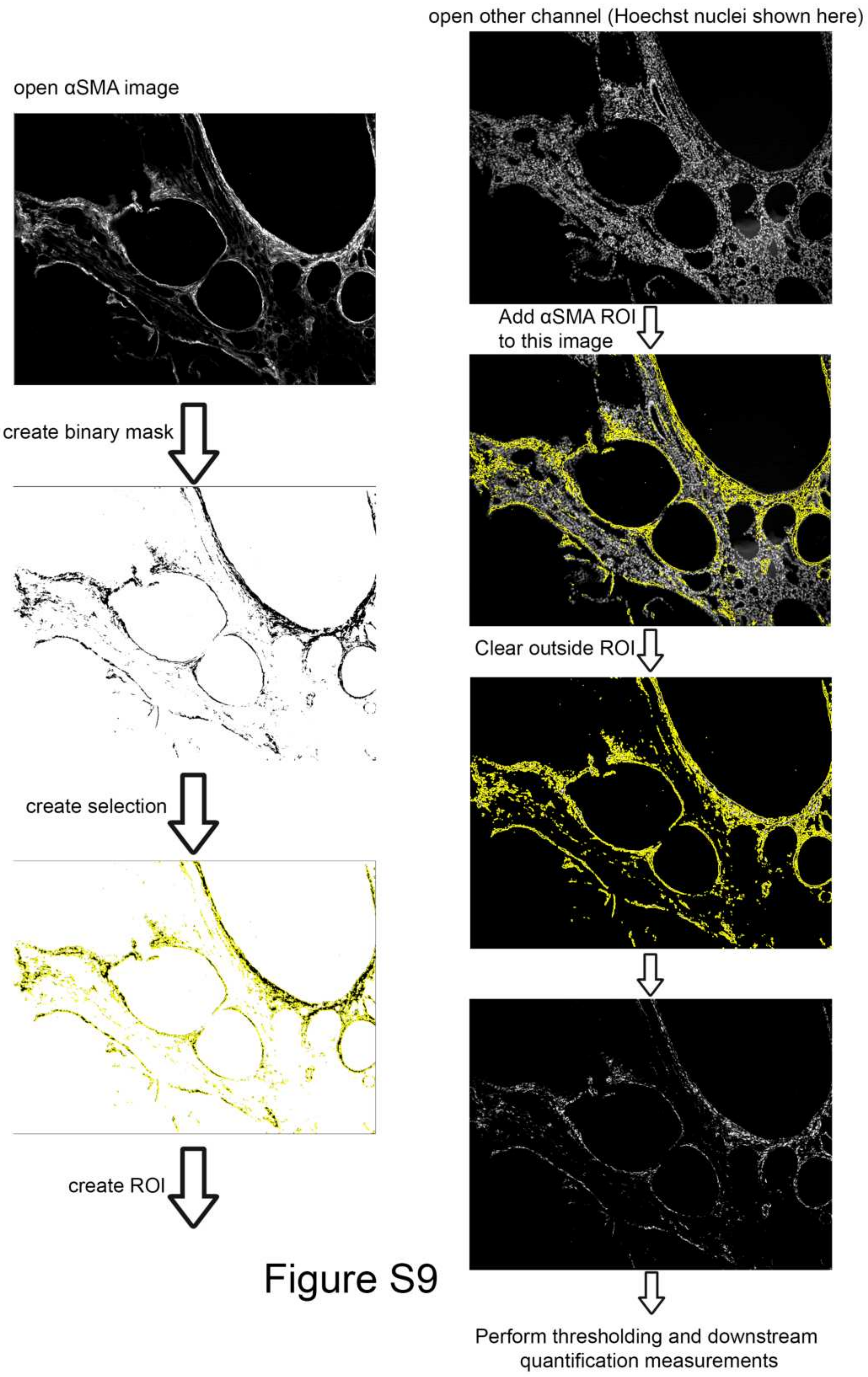
